# Microenvironmental Stiffness Induces Metabolic Reprogramming in Glioblastoma

**DOI:** 10.1101/2023.07.03.547558

**Authors:** Alireza Sohrabi, Austin E. Y. T Lefebvre, Mollie J. Harrison, Michael C. Condro, Talia M. Sanazzaro, Gevick Safarians, Itay Solomon, Soniya Bastola, Shadi Kordbacheh, Nadia Toh, Harley I. Kornblum, Michelle A. Digman, Stephanie K. Seidlits

## Abstract

The mechanical properties of solid tumors influence tumor cell phenotype and ability to invade into surrounding tissues. Using bioengineered scaffolds to provide a matrix microenvironment for patient-derived glioblastoma (GBM) spheroids, this study demonstrates that a soft, brain-like matrix induces GBM cells to shift to a glycolysis-weighted metabolic state which supports invasive behavior. We first show that orthotopic murine GBM tumors are stiffer than peri-tumoral brain tissues, but tumor stiffness is heterogenous where tumor edges are softer than the tumor core. Then, we developed three-dimensional scaffolds with µ-compressive moduli resembling either stiffer, tumor core or softer, peri-tumoral brain tissue. We demonstrate that the softer matrix microenvironment induces a shift in GBM cell metabolism toward glycolysis which manifests in lower proliferation rate and increased migration activities. Finally, we show that these mechanical cues are transduced from the matrix via CD44 and integrin receptors to induce metabolic and phenotypic changes in cancer cells.

## Introduction

Glioblastoma (GBM) is the most common, yet lethal, cancer originating in the brain with a 5-year, overall survival rate of <15%, reported as of 2019^1^. Isolated from peripheral tissue by the blood-brain barrier, GBM tumors rarely metastasize outside of the brain^2^ and instead malignant cells aggressively infiltrate the brain, requiring cancer cells to interact intimately with the unique microenvironment in the brain^3^. Even after maximally safe surgical resection of a primary tumor, diffusely invading GBM cells remain in the peritumoral space. These invading cells display stem-like properties, treatment-resistant phenotypes, and are thought to seed recurrent tumors^4^. The microenvironment surrounding GBM cells includes heterogeneous populations of both cancer and non-cancer cells, vasculature, and extracellular matrix (ECM). While many studies have explored the cellular and soluble components of tumor microenvironment (TME), tumor ECM, a non-soluble factor of TME, has been largely under-appreciated^5^.

Previous studies have reported that the mechanical modulus of the glioma tumor ECM increases with disease progression and have investigated how this increased stiffness may influence GBM cell behavior^6–9^. Increased production of both polysaccharide (e.g., HA) and protein (e.g., fibronectin) components of the ECM leads to elevated stiffness of the tumor ECM^10,11^. Simultaneously, tumor progression correlates with increased expression of ECM receptors, including various integrins and the CD44 receptor for HA, by GBM cells^12,13^. Both integrins and CD44 are able to transduce mechanical cues from the TME into changes in cell phenotype, enabling GBM cells respond to increasing ECM content and stiffness during tumor progression^14,15^.

Our group has demonstrated that patient-derived GBM cells cultured in 3D matrices, in contrast to patient-matched gliomasphere cultures, nimbly acquired resistance to both targeted and conventional chemotherapies^16,17^. Establishing these treatment-resistant culture models required that 3D matrices contain adequate binding sites for both integrins and CD44 receptors and exhibit mechanical properties that approximated peritumoral ECM^18^. Furthermore, we found that invasion of GBM cells through 3D matrices only occurred in these softer, peritumoral-like matrices and required engagement of both CD44 and integrins^18,19^

In the current study, we measured the regional differences in the compressive modulus of the tumor core, tumor edge, and peritumoral ECM tissue in patient-derived GBM tumors orthotopically implanted in mice. We then fabricated ECM-derived matrices with moduli matching these measurements for either tumor core or peritumoral tissue and used these matrices as 3D culture microenvironments for patient-derived GBM cells. Specifically, matrices incorporated tightly controlled amounts of high molecular weight hyaluronic acid (HA) polysaccharides and peptides containing the “RGD” integrin-binding site, which is present in many ECM proteins in the TME^20^. We report that, when cultured in 3D matrices with a modulus approximating the peritumoral ECM, patient-derived GBM cells readily migrated and were able to sense their mechanical environment through the CD44-ezrin signaling axis, which induced a metabolic shift towards increased glycolysis and decreased oxidative phosphorylation. In contrast, when cultured in stiffer matrices with more similar to the tumor core, GBM cells primarily performed oxidative phosphorylation and proliferated, rather than migrated.

Overall, these findings represent a new perspective on the TME, demonstrating that cancer cells can alter their metabolism in response to their local mechanical properties, which in turn determine their behavior. In GBM, this may help to explain why cancer cells at the softer tumor edge readily invade into the softer surrounding parenchyma and rarely metastasize beyond brain into stiffer tissues. Furthermore, we demonstrate that HA-CD44-ezrin has a central role as a mechanosensitive signaling axis that drives these changes. In the future, these results can be applied to develop a better understand the phenotype of invading GBM cells that can be used to identify new therapeutic targets to prevent tumor cell invasion and disease recurrence.

## Results

### GBM tumor is stiffer than peritumoral brain tissue

We performed μ-compression measurements on orthotopically xenografted, patient-derived GBM tumors (HK408 line) using Atomic Force Microscopy (AFM). When animals exhibited set euthanasia criteria (weight loss, motor dysfunction, etc), tissues were harvested, fixed, and vibratome sectioned for measurements. Figure 1.a shows the corresponding Kaplan-Meier survival curves. The μ-compression Young’s moduli of the tumor core (C), tumor edge (E) and the peri-tumoral brain tissue (PT) were measured in each tissue section (Figure 1.b). Measurements of representative sections from different mice, labeled as #1 and #2, are shown in Figure 1.c. Mice #1 and #2 were sacrificed 30 and 35 days after tumor transplantation, respectively. Median Young’s moduli for mice #1 and #2 at different brain locations are reported in Supplementary Table 3 (*p* < 0.0001 for each pairwise comparison). As absolute values of AFM measurements varied across tissue sections, we normalized moduli for each section to the median modulus for the peritumoral region to generate a histogram of normalized median moduli across 2 sections from each of 5 animals (Figure 1.d). Based on this analysis, the average moduli for core (C) and edge (E) regions were 10 ± 7 and 4 ± 3 times stiffer, respectively, than the peritumoral (PT) tissue (Figure 1.d, *p* < 0.0001 for each pairwise comparison).

**Figure 1:**
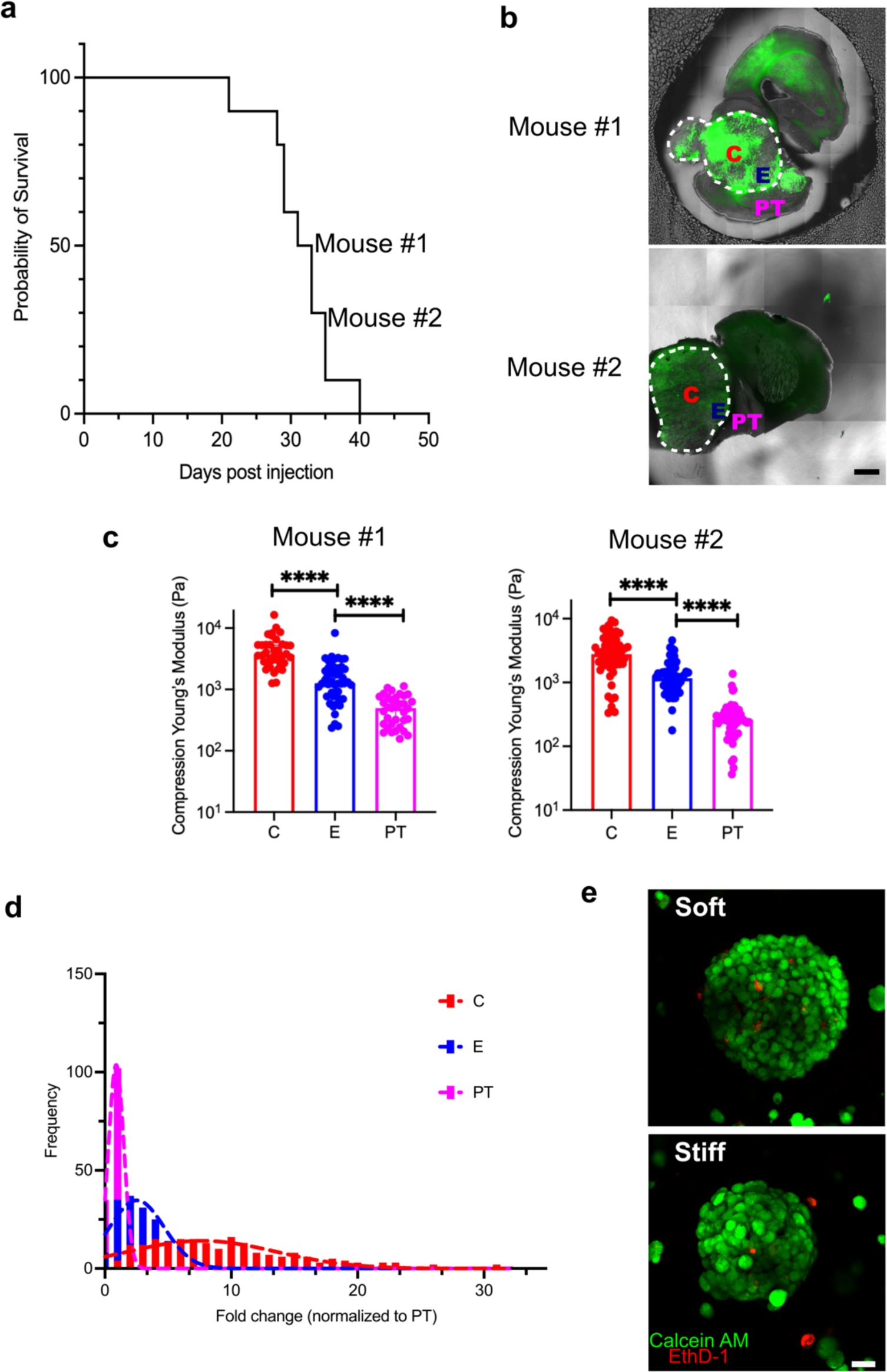
Atomic force microscope (AFM) measurements of μ-compression moduli in GBM xenografts (HK408) explanted from the mouse brain elevated stiffness in tumor, compared to non-tumor, tissue. (a) Kaplan-Meyer survival curve of mice with patient-derived GBM xenografts. (b) Representative tissue sections, collected at the time of euthanasia, of xenografted tumors and their position on the Kaplan-Meyer survival curve. Tumor region is marked with dashed white lines (scale bar = 1mm). (c) Median AFM measurements in explanted tissue slices where “C” indicates tumor core, “E” indicates tumor edge “E,” and “PT” indicates peri-tumoral brain tissue (n = 5 for each region, data showed is the median ± 95% CI, Kruskal-Wallis multiple comparison test, ****: *p* < 0.0001). (d) Histogram of regional AFM measurements of µ-compressive modulus, normalized within each tissue section to measurements of the “PT” region (n=5 mice, Kruskal-Wallis multiple comparison test *p* < 0.0001). (e) Survival of HK177 GBM spheroids cultured within HA hydrogels, 6 days post-encapsulation. Live cells (calcein-AM) are green while dead cells (EthD-III) are red. (Scale bar = 100 μm)

### 3D scaffolds support GBM cells growth

Fabrication of 3D hydrogel matrices is described in detail in our previous work^17,19^ and in the supplementary information. HA hydrogels were fabricated with a wide range of mechanical properties (Supplementary Figure 1.a), specifically to mimic the µ-compressive modulus of either tumor core (stiff hydrogels, E = 3772 ± 1471 Pa) or peritumoral (soft hydrogels, E = 339 ± 87 Pa) ECM (Supplementary Figure 1.b, *p* < 0.0001). Higher stiffness was achieved by increasing crosslinking density, which may affect diffusion of nutrients, growth factors, etc, through 3D culture scaffolds. Thus, we used fluorescence recovery after photobleaching (FRAP) to demonstrate that there were no significant differences in diffusion of large model species through soft and stiff hydrogel matrices (FITC-Dextran, 20 and 70 kDa) (Supplementary Figure 1.c).

GBM spheroids were encapsulated within these hydrogels during crosslinking to establish 3D cultures. All *in vitro* experiments were performed on two unique patient-derived GBM cell lines, HK177 and HK408, the latter of which was used to establish murine xenografts in the study described above. Based on GBM tumor subtype classification system described by Verhaak et al.^21^, HK177 was classified as the mesenchymal-type, and HK408 as the proneural-type^22^. To establish cell-cell interactions, uniformly sized GBM spheroids were formed using Aggrewell^TM^ plates, where spheroid size was determined by the number of cells initially seeded in each micro-well (Supplementary Figure 1.d-h). A single spheroid size, formed from an initial seeding density of 500 cell/μ−well, was used for all following experiments. As results were comparable between HK408 and HK177 cell lines, only data for HK177 cells are shown in the main text, while data for HK408 cells can be found in supplementary materials. Both GBM cell lines exhibited good viability in soft and stiff hydrogels (Figure 1.e, Supplementary Figure 1.i). There were some statistically significant differences (*p* < 0.05) in the percentages of cells undergoing apoptosis in spheroids cultured in soft and stiff hydrogels (Supplementary Figure 1.j). However, differences were inconsistent across lines, where HK177 spheroids were more apoptotic in soft hydrogels (10.1 ± 4.4% vs. 3.9 ± 1.9%, *p* = 0.005) while HK408 spheroids were more apoptotic in stiff hydrogels (5.1 ± 2.5% vs. 8.9 ± 4.4%, *p* = 0.02).

### Soft scaffolds increase GBM cell aerobic glycolysis activity

To investigate stiffness-induced changes in the GBM cell transcriptome, we performed bulk, whole transcriptome sequencing and/or qRT-PCR on RNA extracted from GBM spheroids culture in soft or stiff matrices after 7 days. Across cells cultures in scaffolds with varying stiffnesses, no significant differences in mRNA expression were observed of genes associated with ECM degradation, including matrix metalloproteases (MMPs) and hyaluronidases, or markers associated with proneural or mesenchymal tumor subtypes, including TGFβ1, TIMP1, CHI3L1, and OLIG2 (Supplementary Figure 2.a and b). Likewise, no differences were seen in protein localization of factors that have been widely associated with mechanical signaling, including YAP, whose translocation to the nucleus has been widely associated with a response to a stiff microenvironment (Supplementary Figure 2.c)^23,24^.

Bulk RNA sequencing (RNA-seq) identified that the most differentially expressed transcripts between GBM cells cultured in soft and stiff hydrogels were mitochondrially encoded genes (denoted by “MT-” in the figures), which were consistently down-regulated in soft hydrogels when compared to cultures in stiff hydrogels and gliomasphere culture (GS) (Figure 2.a and Supplementary Figure 3.a). Transcriptomic data were further analyzed using Ingenuity Pathway Analysis (IPA) to identify candidate pathways affected by hydrogel stiffness. For both patient-derived cell lines, oxidative phosphorylation and mitochondrial dysfunction were among the top differentially affected canonical pathways (Figure 2.b and Supplementary Figure 3.b).

**Figure 2:**
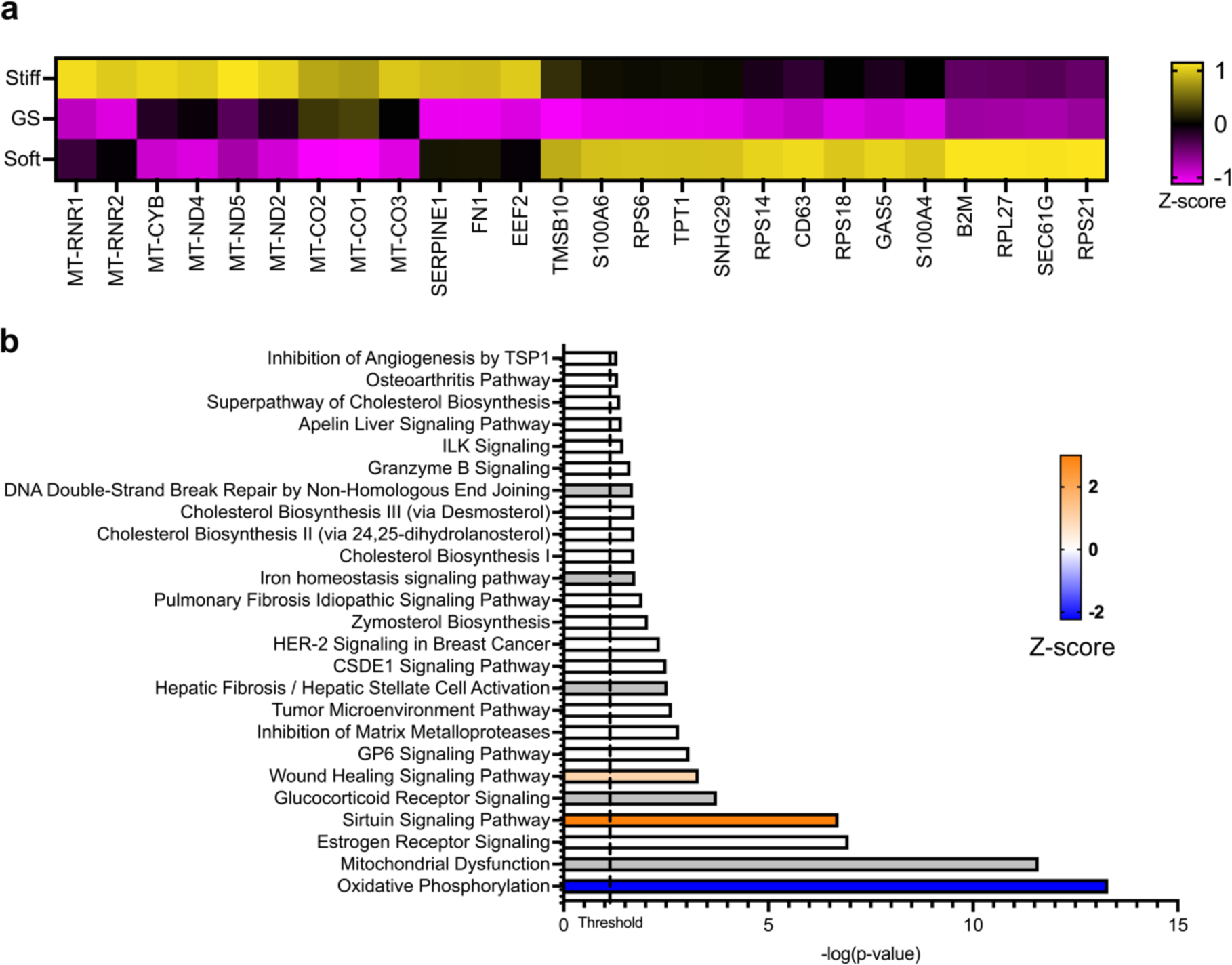
Hydrogel stiffness directly impacts expression of metabolic transcripts. (a) Expression of mitochondrially-encoded genes that participate in the electron transfer chain, denoted with MT-, was lower in soft hydrogel cultures when compared to stiff hydrogel and gliomasphere (GS) cultures. (b) Oxidative phosphorylation and mitochondria dysfunction were the most affected canonical pathways by culture condition, identified using Ingenuity Pathway Analysis (Qiagen). Oxidative phosphorylation was predicted to be lower in soft hydrogels when compared to stiff. Data shown here are for the HK177 GBM cell line.

To investigate whether these transcriptomic differences reflected metabolic changes in GBM cells, we performed fluorescence lifetime imaging (FLIM) of NAD^+^ and NADH, which can be distinguished because bound NADH has a higher fluorescence lifetime (3.4 ns) than free NAD^+^ (0.4 ns) ^25^. Increased presence of free NAD+ in the cytosol generally indicates an increase in GLY, as this NAD+ is not being shuttled to mitochondria to fuel OXPHOS^25^. Thus, the ratio of bound NADH to free NAD+ is used to estimate the relative activities of OXPHOS and GLY in a cell, where a larger ratio indicates higher OXPHOS compared to GLY activity^26^. In our investigation, GBM cells cultured in soft hydrogels had a lower fraction of bound NADH than those cultured in stiff hydrogels or as free-floating spheroids (Figure 3.a and b, Supplementary Figure 4.a and b), indicating that cells in soft hydrogels rely more on glycolysis than OXPHOS while those in stiff hydrogels or gliomaspheres rely more on OXPHOS than glycolysis^27,28^. Within each biological replicate, figures show the fraction of bound NADH normalized to its median value for cells in stiff hydrogels.

**Figure 3:**
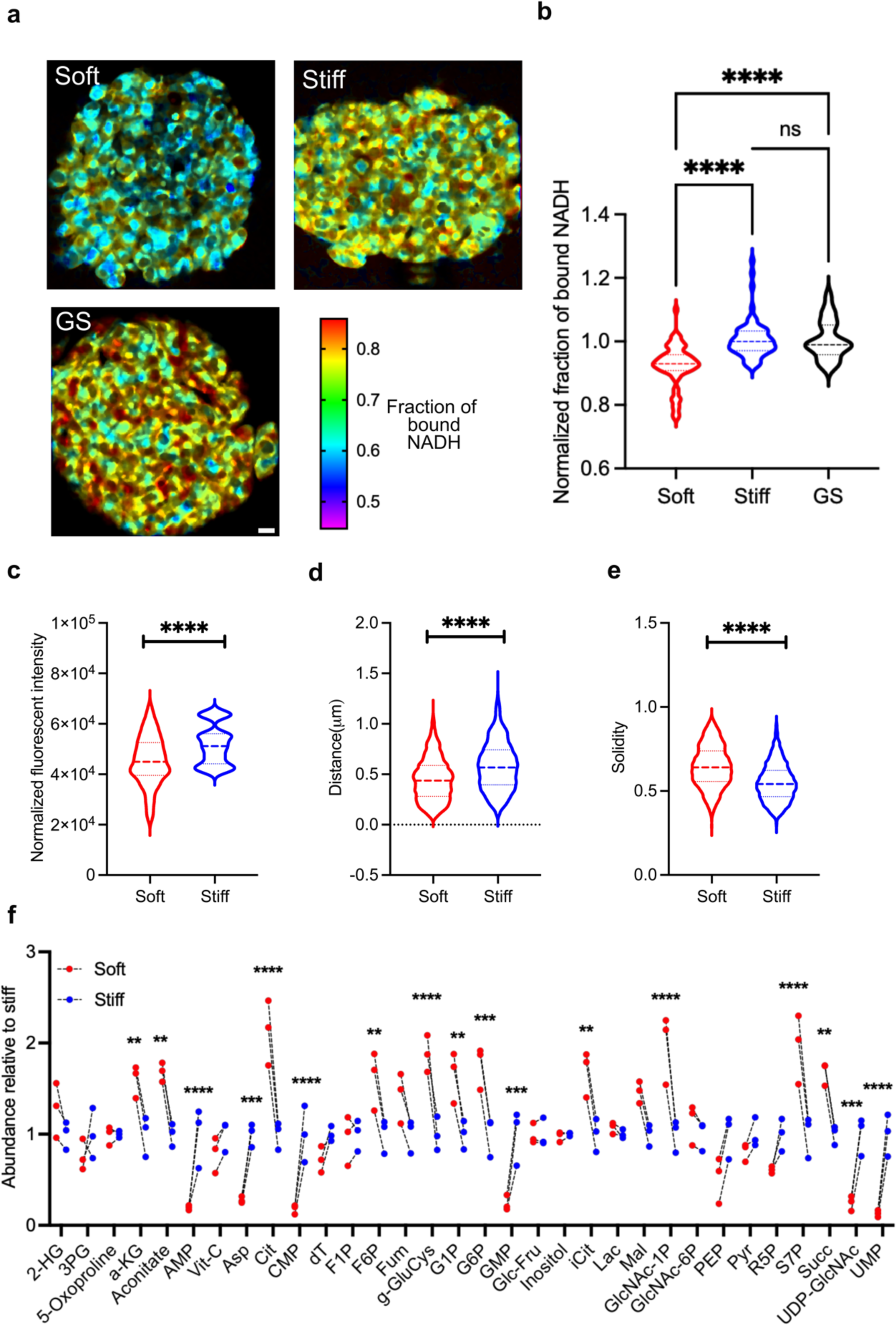
GBM cells cultured within soft hydrogels underwent a metabolic shift toward dominance of aerobic glycolysis (GLY). (a) Representative NADH fluorescence lifetime imaging of GBM cells cultured in 3D hydrogels or suspended gliomasphere (GS) (scale bar = 20 μm). (b) Bound fraction of NADH decreases in soft hydrogel cultures, suggesting higher GLY activity (n = 3 biological repeats, 9 GBM spheroids for each condition in each biological repeat, Mann-Whitney non-parametric test, *\*\*\*\*: p* < 0.0001). (c) Mitochondria content analysis revealed fewer mitochondria, (d) lower distance traveled by mitochondria tracks, and (e) higher solidity of mitochondria tracks (i.e., less elongation) in soft hydrogel cultures (n = 5, Mann-Whitney non-parametric *\*\*\*\*: p* < 0.0001)(f) Abundance of released metabolites: TCA cycle intermediate metabolites were more abundant in soft hydrogels. On the other hand, nucleotide monomers abundance were higher in stiff hydrogels. (n = 3, unpaired, two-sided t-test, *: *p* < 0.05, **: *p* < 0.01, ***: *p* < 0.001, ****: *p* < 0.0001). Data shown here are for the HK177 GBM cell line.

As oxygen availability can directly affect cell metabolism, we investigated the oxygen availability in GBM spheroids cultured in soft or stiff scaffolds using the Image-iT^TM^ hypoxia dye, which becomes fluorescent, and remains fluorescent, if at any time when the local oxygen content falls below 5%. No hypoxia was observed after 1 or 3 days in culture in any scaffold conditions. By day 6, green fluorescence from Image-iT^TM^ hypoxia dye appeared and was more pronounced near the centers of spheroids encapsulated within stiff hydrogels (Supplementary Figure 5). However, we did not observe expression of the hypoxia-induced factor-1α (HIF-1α) protein in any conditions during the experimental period (Supplementary Figure 6). Together, these data strongly indicate that matrix stiffness affects GBM cell metabolism independently of the effects of hypoxia.

Mitochondrial numbers and network morphology, which can be used to predict a cell’s metabolic state^29,30^, were characterized from mean fluorescent intensity (MFI) of a Mito Tracker probe (Supplementary Video 1). In soft hydrogels, HK177 and HK408 cells showed approximately 20% and 60%, respectively, lower MFI (Mito Tracker) than in stiff hydrogels, indicating the presence of fewer mitochondria (Figure 3.c, Supplementary Figure 4.c, *p* < 0.0001). Morphology of mitochondrial networks was analyzed using distance and solidity parameters, as previously described^30^, where a more continual and elongated mitochondrial network indicate increased OXPHOS activity^31–34^.

The median of the distance parameter, summations of the linear distance across all segments of each mitochondrial track, was consistently lower for both HK177 and HK408 cells when cultured in soft, compared to stiff, matrices (Figure 3.d, Supplementary Figure 4.d *p* < 0.0001). Likewise, the median of the solidity parameter, which measures the roundness of an object where a value of 1 is completely circular and a value of 0 indicates a fully elongated object^30^, was higher for cells in soft hydrogels (Figure 3.e, Supplementary Figure 4.e, *p* < 0.0001). Together, these measurements indicate that GBM cells cultured in soft matrices relied less on OXPHOS than those cultured in stiff matrices. We then assessed the mitochondrial networks in patient-derived, orthotopic murine xenografts from immunofluorescence images of MT-CO1 protein, which is also known as complex 1 in the electron transport chain. Mitochondrial networks *in vivo* appeared less elongated near the relatively softer tumor edges, indicating a glycolysis-weighted metabolism, and more elongated near the stiffer tumor core, indicating an OXPHOS-weighted metabolism (Supplementary Figure 7).

To further assess metabolic differences in GBM cells cultured in soft and stiff 3D matrices, we profiled secreted metabolites using mass spectroscopy. Medium collected from GBM cells cultured in soft hydrogels contained a higher abundance of the tricarboxylic acid (TCA) cycle intermediates than medium collected from cultures in stiff hydrogels, indicating that these species are being secreted rather than being consumed during OXPHOS^35^. For example, cultures in soft hydrogels had higher relative abundances of TCA cycle intermediates aconitate (*p* = 0.0013), succinate (*p* = 0.0015), citrate (*p* < 0.0001), iso-citrate (*p* = 0.0011), and ⍺-ketoglutarate (*p* = 0.0097) (Figure 3.f). Functionally, the higher abundance of TCA intermediates in collected medium from soft hydrogel cultures indicates a reduction in TCA flux and potentially lower OXPHOS activity^35^.

In contrast, the relative abundance of nucleotide monomers, including uridine monophosphate (UMP) (*p* < 0.0001), guanosine monophosphate (GMP) (*p* = 0.0002), adenosine monophosphate (AMP) (*p* < 0.0001), and cytidine monophosphate (CMP) (*p* < 0.0001), was higher in medium collected from stiff hydrogel cultures (Figure 3.f). The higher abundance of nucleotide monomers in medium from in stiff hydrogel cultures suggests a more proliferative activity^36^.

### Stiffness-induced metabolic shift alters GBM proliferation and migration

In line with previous studies found that higher OXPHOS activity is necessary for cell proliferation^31,37,38^, we report increased proliferation rates for GBM cells cultured in stiff hydrogels, which exhibited dominant OXPHOS activity, compared to those in soft environments. Within a 3-hour period, 9.3 ± 0.8% and 5.3 ± 1.0% of HK177 cells, cultured in stiff and soft hydrogels, respectively, had proliferated (Figure 4.a, *p* < 0.0001). While less proliferative overall, HK408 cells also proliferated faster in stiff hydrogels, with 3 ± 0.2% and 1.8 ± 0.1% of cells having proliferated within 3 hours in stiff and soft hydrogel cultures, respectively (Supplementary Figure 8.a, *p* = 0.001). In agreement with several previous reports that cancer cell motility increases with higher glycolytic activity^39^, GBM cells had extensive migratory behavior in soft hydrogels, yet did not migrate in stiff hydrogels (Supplementary Figure 9.a and b). Furthermore, while GBM cells within spheroids relied more on GLY in soft hydrogels, cells migrating out from the spheroid into the surrounding matrix exhibited a metabolic shift toward OXPHOS compared to their parental spheroids (Supplementary Figure 10, *p <* 0.0001 for HK177 and *p* = 0.0036 for HK408),

**Figure 4:**
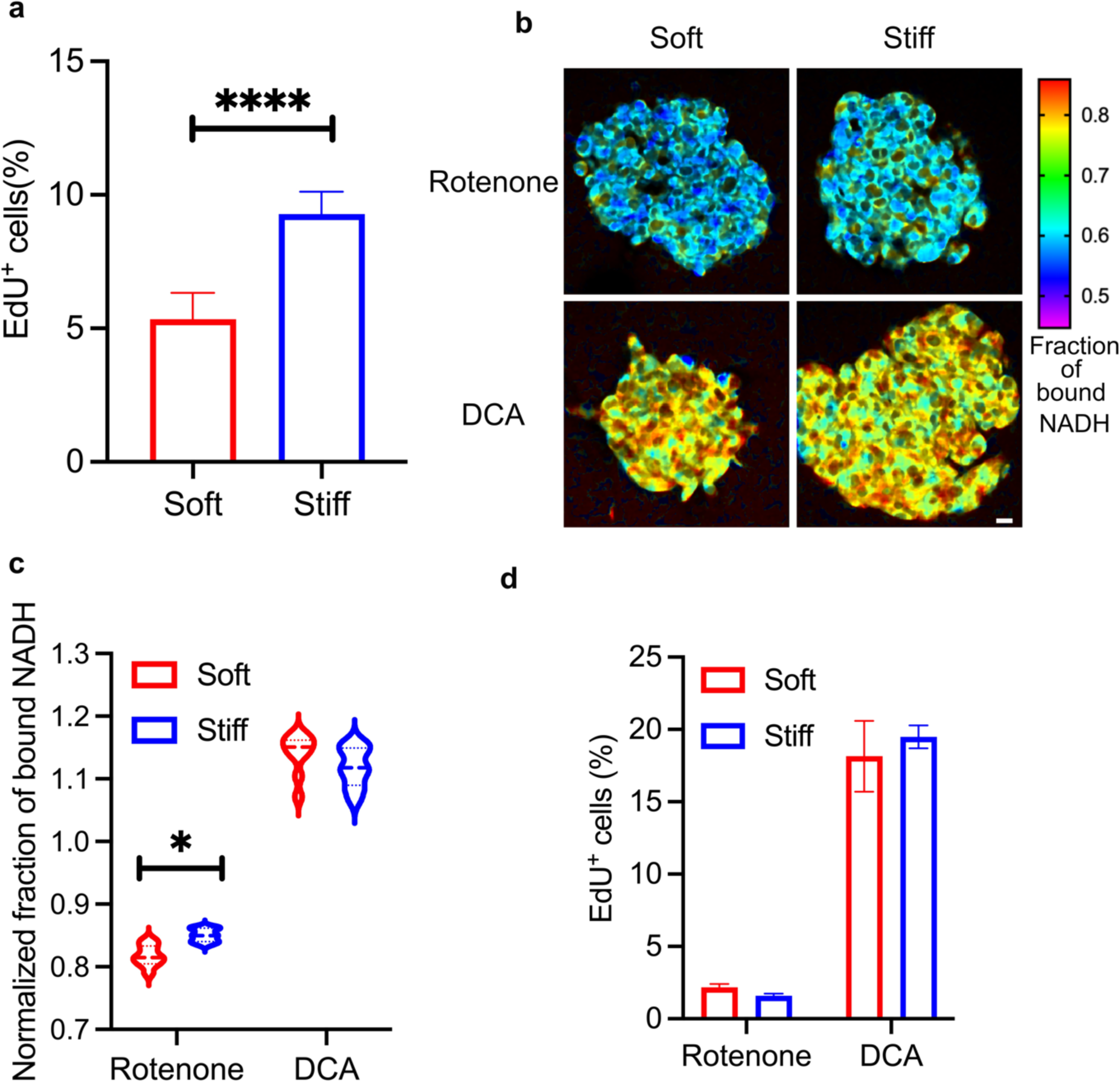
GBM spheroids in soft hydrogels exhibited increased glycolytic activity leaded to decreased proliferation. (a) An EdU incorporation assay found lower proliferation rates for cells cultures in soft hydrogels when compared to those in stiff hydrogels (n = 2 independent repeats, each with 3 culture replicates, reported values are Mean ± SD, Welch’s t-test *\*\*\*\*: p* < 0.0001). (b) Representative FLIM images of GBM cells treated by metabolic-altering small molecules (scale bar = 20 μm) and (c) Normalized fraction of bound NADH confirming the ability of rotenone and DCA to inhibit OXPHOS and GLY, respectively (n=2 independent repeats, 2-way ANOVA and Tukey multiple comparison, *: *p* = 0.0271). (d) In both culture environments (stiff and soft hydrogels), inhibition of GLY using DCA resulted in increased proliferation rates (EdU assay) while inhibition of OXPHOS using rotenone treatment decreased proliferation rates. No differences between culture environments were observed (n = 2 biological replicates, 3 individual measurements for each condition in each biological repeat, Mean ± SD, 2-way ANOVA multiple comparison, rotenone soft vs stiff *p* = 0.8713, DCA soft vs stiff *p* = 0.3102). Data shown here are for the HK177 GBM cell line.

We further investigated the relationship between proliferation rate and metabolic activity using rotenone and dichloroacetate (DCA), which inhibit the OXPHOS and glycolysis pathways, respectively^40,41^. First, FLIM measurements confirmed the effectiveness of rotenone and DCA in altering the expected metabolic pathways in 3D GBM cell cultures (Figure 4.b and c, Supplementary Figure 8.b and c). For both HK177 and HK408 cell lines, inhibition of OXPHOS using rotenone resulted in decreased proliferation, while inhibition of glycolysis using DCA increased proliferation. Specifically, rotenone decreased the percentages of HK177 cells which had proliferated during a 3-hour period to 2.2 ± 0.2% and 1.6 ± 0.1% in soft and stiff hydrogels, respectively (compared to respective non-treated conditions, *p* < 0.0001) (Figure 4.a and d). For the same HK177 cell line, DCA increased percentages of proliferating cells to 18.2 ± 2.5% and 19.5 ± 0.8% in soft and stiff hydrogels, respectively (Figure 4.a and d, compared to respective non-treated condition *p* < 0.0001). Similarly, rotenone decreased percentages of HK408 cells proliferating within 3 hours to 0.0 ± 0.0% and 0.1±0.0% in soft and stiff hydrogels, respectively, (compared to the non-treated conditions, *p* = 0.0008 for soft and *p* < 0.0001 for stiff) and DCA increased proliferation percentages to 4.0 ± 0.5% and 3.4 ± 0.3% in soft and stiff hydrogels, respectively (Supplementary Figure 8.a and d, compared to the respective non-treated conditions *p* = 0.0174 for soft and *p* = 0.221 for stiff).

### CD44 and integrins mediate stiffness signaling

Tumor cells interact with their surrounding ECM through cell receptors, including such RGD-binding integrins and HA-binding CD44^3^, both of which can transduce mechanical cues^6,14^. To investigate whether integrins or CD44 mediated the stiffness-induced metabolic shift of GBM cells cultured in soft hydrogels, we used the small molecule inhibitors Cilengitide^TM^ (CRGD) and NSC668394. CRGD, a cyclic peptide containing RGD, is a competitive inhibitor of integrin-RGD binding in the hydrogel matrices^18,42^. NSC668394 inhibits phosphorylation of ezrin, an adaptor protein of the ezrin/radixin/moesin (ERM) subfamily, whose phosphorylation by CD44 or various integrins mediates cell migration^43,44^. Immunofluorescence images show overlapping expression of CD44 and ezrin at cell membranes, indicating that these receptors are present to phosphorylate ezrin and possibly convey information about the 3D matrix to GBM cells (Supplementary Figure 11).

FLIM results showed that CRGD had minimal effects on the metabolism of HK177 cells in 3D cultures. While cells in both soft and stiff hydrogel cultures experienced an overall increase in relative OXPHOS activity, glycolysis still dominated metabolic activities in soft hydrogel cultures (Figure 5.a and b, *p* < 0.0001). In contrast, ezrin inhibition induced a larger increase in OXPHOS activity in all conditions and eliminated metabolic differences between soft and stiff hydrogel cultures (Figure 5.a and b, *p* = 0.8076). These data indicate that the shift to glycolysis dominance in softer microenvironments is mediated by CD44-ezrin interactions in the HK177 cell line. However, for HK408 cells, both CRGD treatment and ezrin inhibition increased OXPHOS and eliminated metabolic differences between soft and stiff hydrogel cultures (Figure 5.a and c, *p* = 0.2799 for CRGD comparison, *p* = 0.9037 for ezrin inhibition comparison). These results indicate that in the HK408 cell line ezrin phosphorylation and integrin engagement both affect metabolic changes induced by the mechanical microenvironment.

**Figure 5:**
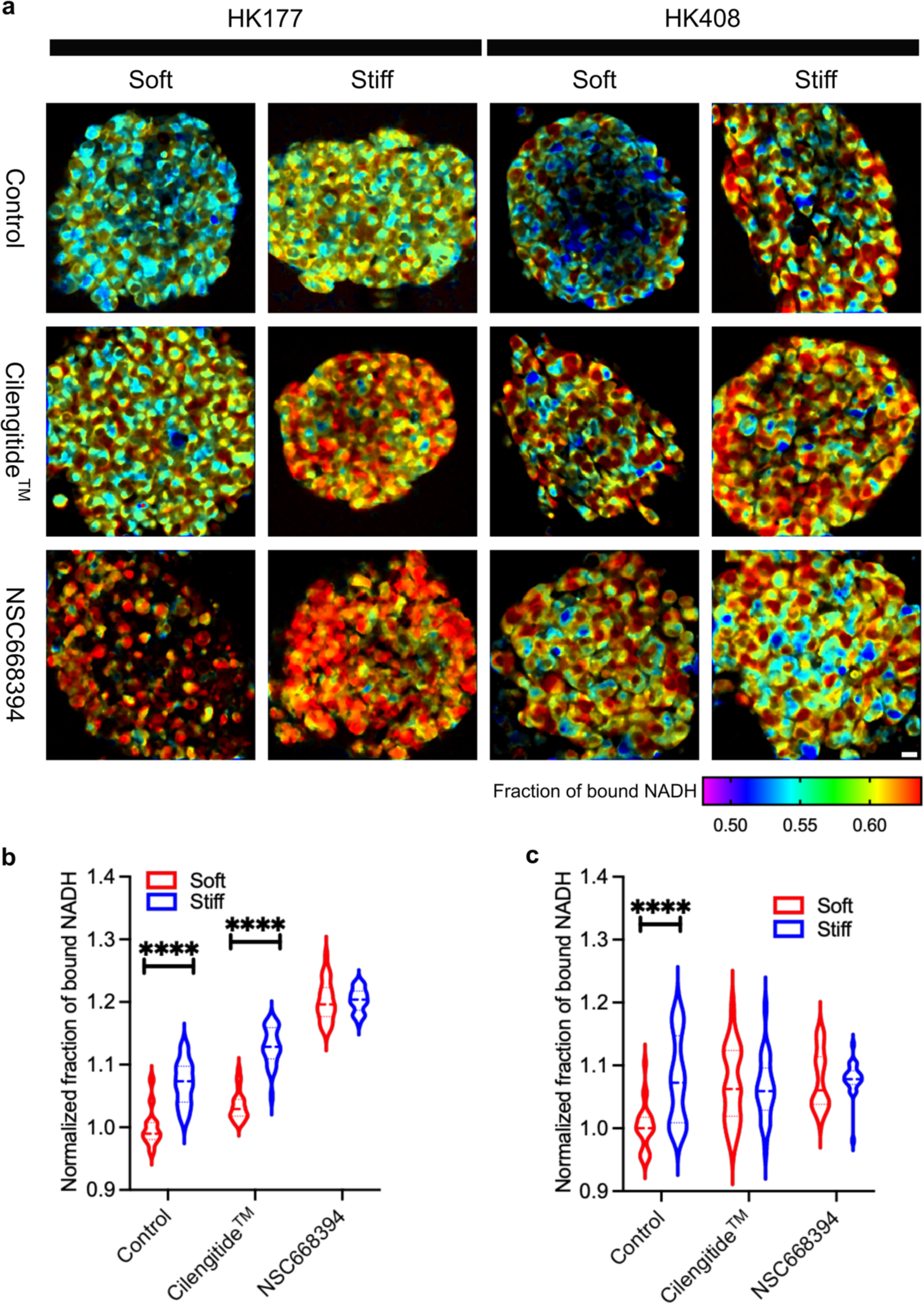
CD44 and integrins are involved in transmitting information about the mechanical properties of the surrounding matrix. (a) Representative FLIM images of HK177 and HK408 GBM cell cultures when treated with Cilengitide^TM^ (RGD-binding inhibitor) and/or NSC668394 (ezrin inhibitor) (scale bar = 20 μm). (b) For the HK177 cell line, the normalized fraction of bound NADH shows that only inhibition of HA/CD44 interaction, using NSC668394, eliminated metabolic differences between GBM cells cultured in soft and stiff hydrogels (n = 3 biological repeats, 9 GBM spheroids for each condition in each biological repeat, two-way ANOVA and Tukey multiple comparison, ****: *p* < 0.0001). (c) For the HK408 cell line, both CD44/HA, using NSC668394, and integrin/RGD (using Cilengitide^TM^) interactions were important for the metabolic shifts (n = 3 biological repeats, 9 GBM spheroids for each condition in each biological repeat, two-way ANOVA and Tukey multiple comparison, ****: *p* < 0.0001).

## Discussion

The TME has a crucial influence on GBM progression and recurrence^3^. Among features of the TME, mechanical stresses have been reported to impact tumor progression^6,8,9^. In this investigation, we used a 3D culture system to isolate the effects matrix elasticity from other features of the TME, including ECM composition and density, to discover that the ECM stiffness directly induces shifts in metabolism, which in turn influences GBM cell behavior.

We report that patient-derived, orthotopically xenografted GBM tumors in mice consistently exhibit μ-compression moduli 3-17 times larger than that of the tumor-adjacent brain tissue. This finding is in agreement with previous studies reporting that GBM tumor tissue is stiffer (i.e., has a higher compressive, elastic modulus) than normal brain tissue^7,45^. Increased tissue elasticity has been attributed to increased net deposition of ECM by GBM and other cells in the TME^10^. As the goal of these studies was to investigate the specific influences of elasticity on tumor cells, we focused on the μ-compression modulus of the tissue matrix. However, it should be noted that, *in vivo*, additional mechanical changes, such as increases in cerebrospinal fluid pressure are well-known to occur^46,47^ and likely also contribute to GBM cell behavior but were not evaluated within the scope of this report. Additionally, while formation of necrotic cores is uncommon in murine xenografts, GBM tumors in patients often have necrotic cores, which are largely composed of fluid. Thus, we expect that the core area measured in xenografts is analogous to the tumor area between the necrotic core and the invasive edge in patients.

We posit that the heterogeneous mechanical landscape of tumors may contribute to phenotype differences between GBM cells residing in the stiffer core and softer invasive edge of the tumor. For example, while a mixture of cells exhibiting proneural or mesenchymal subtypes can be found in both the edge and core of GBM tumors, the edge cells have a higher capacity for invasion^48–50^ and distinct transcriptional states from the core cells^48,51^. Furthermore, it has been proposed that GBM recurrence relies on edge cell plasticity, where these are thought to transition to a more core-like state immediately prior to seeding a new tumor^48,49^. In these studies, we provide additional evidence that spatial heterogeneity in ECM stiffness contributes to regional differences in GBM cell phenotype and plasticity by altering metabolism, where the relatively soft tissue surrounding tumors promotes a transition to an invasive state.

In contrast to previous reports, we did not find any significant differences in mRNA expression of genes whose expression have been commonly associated with ECM degradation, including matrix metalloproteases (MMPs) and hyaluronidases, or proneural or mesenchymal GBM subtypes, including TGFβ1, TIMP1, CHI3L1, and OLIG2. Likewise, although previous studies have associated CD133 as a marker of edge cells and CD109 as a marker of core cells in GBM tumors^48–50^, we did not observe differences in CD133 or CD109 expression in GBM cells culture in scaffolds mechanically matched to the tumor edge or core. Finally, no differences were seen in expression or subcellular localization of proteins that have been widely associated with mechanical signaling, including YAP, whose translocation to the nucleus has been widely associated with a response to a stiff microenvironment^52,53^. While YAP may still be involved in other cellular functions, such as sequestering β-catenin^54^, YAP nuclear localization is more complex in 3D hydrogels and may require a stiffer ECM than explored in this study^24,55,56^.

Transcriptomic analysis on the 7^th^ day of culture did indicate decreased expression of mitochondrially encoded genes, in particular those involved in electron transport chain, by GBM cells cultured in soft hydrogels compared cells in stiff hydrogels or gliomaspheres. This finding suggested a metabolic shift towards glycolysis and was supported by further investigation using FLIM (single-cell analysis), analysis of mitochondrial network structure, and LC-MS-based metabolomics (population analysis). While we see clear differences in metabolic activity for GBM cells cultured within soft, tumor edge-like scaffolds at 7 days, downstream, metabolism-induced changes in gene expression – for example, related to a shift in proneural or mesenchymal phenotype – may take longer to occur.

Furthermore, we provide evidence of the potential *in vivo* relevance of these *in vitro* results, obtained using our bioengineered scaffold cultures, by investigating the mitochondria network structure in patient-derived, orthotopic xenograft tissues. Similar to soft hydrogel cultures, GBM cells at the softer edges of xenografted tumors had more rounded and fragmented mitochondrial networks. In contrast, like stiff hydrogel cultures, GBM cells in stiffer tumor core had more elongated mitochondrial network. However, differences in mitochondrial network structure were less pronounced in tumor xenografts than in GBM 3D cultures, perhaps indicative of the more heterogenous micro-scale stiffness landscape in xenografts, compared to hydrogels and the presence of additional mechanical forces (e.g., fluid pressure) and microenvironmental cues not captured in the *in vitro* model.

The TME is deeply intertwined with metabolism, where metabolic products condition the interstitial space. Alterations in metabolism are common in cancer cells and typically occur as precursors to changes that require longer time scales (e.g. transcriptomic, proteomic, and phenotypic changes). We speculate that the stiffness-induced metabolic shift observed in GBM cells reflects an early stage of tumor development and potentially can be used to predict downstream behavioral changes^57^. While previous research investigating cancer cell metabolism has often focused on relationships to oxygen availability^58^, our results suggest the existence of an initial, hypoxia-independent mechanism by which GBM cell metabolism is directly influenced by ECM stiffness. Using a reagent that becomes fluorescent when local oxygen levels fall below 5%, we confirmed that oxygen levels in 3D cultures predominately remained above 5%, with the exception of larger spheroids in stiff hydrogel cultures which began to experience lower oxygen levels in spheroid centers around day 6 in culture.

The lower available oxygen in spheroids cultured in stiff hydrogels may be attributed to higher proliferation rates, leading to larger spheroids in which diffusion to the spheroid center may be limited. In a handful of individual large cultured in stiff hydrogels, FLIM analysis indicates that GBM cells in the center had increased levels of glycolysis by day 7 in culture. Regardless of the cause, HIF1-α expression, which is commonly used as an indicator that a cell has experienced sustained hypoxia, was not observed in any conditions evaluated here. While longer culture times may lead to hypoxia-induced metabolic changes, at the shorter culture times assessed here the metabolic shift towards glycolysis in soft hydrogel cultures appears to be independent of oxygen availability.

Changes in metabolic signatures directly affect cancer cells behavior. In agreement with previous reports^39,59^, increased glycolytic activity in soft hydrogel cultures of GBM spheroids correlated with a higher propensity for invasion. *In vivo*, tumor edge cells, residing in softer microenvironment, are likewise highly invasive^48–50^. In contrast, the OXPHOS-weighted metabolism observed in stiff hydrogel cultures correlated with a higher proliferation rate, similar to the hyperproliferative, pseudopalisading regions typically observed near the GBM tumor core, just outside of the necrotic area in patients^48^. This result aligns with the idea that cells, energetically, have the capacity to either “go-or-grow”^60,61^. Previous studies have indicated that migration through a denser matrix, like the stiff hydrogels used here, requires more energy^62,63^, a situation in which cells may instead only have sufficient ATP available to proliferate. Decision to migrate or proliferate is governed based on ECM cues. In soft hydrogel cultures, where cells require less energy to migrate than in denser matrices, GBM cells may initiate migration by switching to a reliance on glycolysis to produce more ATP faster. However, as GBM cells invading soft, 3D hydrogels and moved away for their parental spheroid, they reverted to an OXPHOS-dominant metabolism. This shift may be necessary to sustain migratory activity, which demands consistent energy production^64^.

GBM tumor cell phenotypes and treatment responses vary greatly both within a single tumor and across patients. Our data suggests that this heterogeneity can be at least partly attributed to the ECM content and stiffness. The CD44 receptor for the HA polysaccharide and integrin receptors for various ECM proteins have each been reported to relay mechanical cues from the matrix to cells^6,14^. Here, we showed that both CD44 and integrins on GBM cells can transduce mechanical cues that mediate downstream metabolic shifts, but the extent of their influence depends on the patient-derived cell line. For example, FLIM measurements in HK408 cultures had a very large distribution demonstrating a heterogeneous response across the cell population. In contrast, the HK177 cell population had a more uniform response.

While stiffness-induced phenotypic shifts along the proneural-to-mesenchymal axis or in stemness were not observed from transcriptomic analysis, there were noteworthy changes in mRNA expression of genes associated with cell invasion and tumor stiffening in cancers. First, several ribosomal genes (RPS6, RPS14, RPS18, RPS21, RPL27), previously found to be upregulated with increased cancer cell migration, were overexpressed by cells cultured in soft hydrogels^65,66^. In both esophageal squamous cell carcinoma and ovarian cancer cells, RPS6 has been reported to induce overexpression of matrix-degrading enzymes, including MMP2 and MMP9^67,68^, which facilitate cell migration. Second, calcium-binding protein subunits (S100A6, S100A4), previously correlated with worse clinical prognosis in colorectal^7,69^ and breast cancers^70^, were overexpressed by the migratory GBM cell populations in soft hydrogel cultures. Third, increased expression of CD63, a marker of exosomes, was observed in soft hydrogel cultures, possibly allowing GBM cells to more closely communicate with the stromal cells near the tumor border^71^. Finally, GAS5, a non-coding long RNA, was upregulated in soft hydrogel cultures. While GAS5 has been found to be a tumor suppressor, it also promotes autophagy, which could in turn facilitate metabolic processes in cancer cells to promote tumor growth^72,73^.

In stiff hydrogels cultures, GBM cells overexpressed genes associated with ECM remodeling and stiffening (i.e., SERPINE1 and FN1), indicating a possible positive feedback loop where GBM cells exposed to a stiffer matrix in turn act to further stiffen the TME. SERPINE1 encodes for expression of plasminogen activator inhibitor 1 (PAI-1), which is a physiological inhibitor of serine proteases, urokinase-type plasminogen activator, and tissue-type plasminogen activator^74^. Inhibition of these proteins by PAI-1 results in inhibition of plasminogen-to-plasmin conversion as well as plasmin-dependent activation of MMPs^75^. Thus, higher expression of SERPINE1 in stiff microenvironment, could potentially increase the concentration of PAI-1 which in turn inhibits ECM degradation and protects ECM proteins from proteolytic degradation^76,77^. Inhibition of ECM degradation, in addition to higher expression of fibronectin (FN1) and other ECM components would be expected to effectively stiffen the TME.

In sum, this work describes how GBM cells can relay mechanical cues from their microenvironment to rewire their metabolism to induce either proliferative or invasive behaviors. In particular, results indicate that a softer matrix, similar to at the border of GBM tumors *in vivo*, promotes a metabolic shift towards increased glycolysis and invasive activity. In contrast, GBM cells residing in a stiffer microenvironment, with mechanical properties resembling the tumor core, or in suspension-cultured spheroids supports an OXPHOS-dominant metabolism and increased proliferation. These results provide evidence that the mechanical microenvironment at the tumor-brain tissue interface may drive GBM cells at the tumor edge to adopt an invasive behavior characteristic of GBM tumors. These studies contribute to a better understanding of the relationship between ECM mechanics, cell metabolism, and tumor progression that we anticipate will pave the way for development of new, targeted therapies for GBM.

## Methods

### Tumor xenografts

Mice were prepared for aseptic surgery in accordance with protocols set by UCLA’s Division of Laboratory Animal Medicine. HK408 cells (proneural), constitutively expressing firefly luciferase and green fluorescent protein (GFP), were dissociated with TrypLE (Thermo Fisher) and resuspended to 10^5^ cells per 3 μL in growth medium. 3 μl of cell suspension was implanted into the right striatum of NSG mice at 0.5 mm anterior and 1.0 mm lateral of bregma, and 2.5 mm deep. Tumor growth was monitored by luminescence imaging on an IVIS Illumina II system at the Crump Institute’s Preclinical Imaging Technology Center. Mice brains were fixed in 4% PFA overnight and then sectioned into 100 µm slices using a Leica VT1200S Vibratome. Brain slices were adhered to glass slides using Cell-Tak^TM^ cell and tissue adhesive (Corning)^78^.

### Atomic force microscopy (AFM)

All AFM measurements were done on a Bruker Nano wizard 4 atomic force microscope at Nano/Pico Center (NPC), California Nano System Institute (CNSI). We used silicon nitride cantilevers equipped with 2.5 µm (diameter) silicon dioxide particles (NovaScan, nominal spring constant of 0.01 N/m). AFM measurements were carried out in PBS at 37 °C. Post-installation, the probe was allowed to thermally equilibrate in the PBS buffer for 1 hour. The cantilever sensitivity was measured using a generated force-curve on the glass slide. In addition, probe’s spring constant was measured using the manual thermal calibration. Measurements were done at 0.2 μm/s indentation speed with a total indentation of 1 μm. We measured a matrix of 32 X 32 μm with 4 μm interval (8 X 8 grid) in tumor core, edge and the peri-tumoral tissue. Data analysis was carried out in JPKSPM data processing software using the Snodden model for spherical probes. Data points correspondent to “no-contact” spots were removed from final analysis.

### Cell culture

Patient-derived GBM cells, HK177 (mesenchymal) and HK408 (proneural) (Both IDH wildtype), were generously provided by Dr. Harley Kornblum at UCLA. All cell lines were collected with strict adherence to UCLA Institutional Review Board protocol 10-000655. We examined cell cultures routinely for negative presence of mycoplasma contamination (Life Technologies, C7028). GBM cells (100,000 cells/mL) were cultured in DMEM/F12 with 1xG21 (Gemini Bio), 1 v/v% normocin (Invivogen), 50 ng/ml EGF (Peprotech), 20 ng/mL FGF-2 (Peprotech), and 5 μg/mL heparin (Sigma-Aldrich). For encapsulation, 24-hour after spheroid formation in Aggrewell^TM^, spheroids were harvested from the wells, centrifuged briefly (200G, 1 min) and resuspended in the hydrogel precursor solution. Hydrogels were formed as described in supplementary information. Post-formation, spheroid-laden hydrogels were transferred to 24-well well plates and cultured in 500 μl of GBM media. Half of the media volume (250 μl) was replaced by fresh media every other day.

### Survival assay

Live/dead assay was performed using LIVE/DEAD^TM^ viability/cytotoxicity kit for mammalian cells (ThermoFisher Scientific) following the manufacturer protocol. The staining solution was prepared in PBS. Hydrogels were incubated with the staining solution for 45 minutes in the culture incubator. Post-staining, hydrogels were washed in PBS before imaging. Images were taken using a Leica SP5 confocal microscope at Advanced Light Microscopy and Spectroscopy lab (ALMS) at UCLA CNSI. Images were taken using a 10X objective. To capture each sphere, Z-stacks with 6 μm were used. Finally, Z-stacks were transformed to 2D images using the Max-projection in ImageJ.

### GBM cell proliferation

GBM cell proliferation was measured using EdU assay (Cayman chemical company). Stock solution of EdU was prepared at 10mM in DMSO and stored in −20°C. For experiments, EdU solution was prepared at 10 μM in GBM media. GBM spheroids-laden hydrogels were pulsed with EdU-containing media for 4 hours in the culture incubator. Post-incubation, excess EdU was washed with PBS containing 1% BSA. GBM cells were extracted using an established protocol^79^. Post-extraction, GBM cells were fixed in 4% PFA for 15 minutes at room temperature followed by washing with PBS-1% BSA. GBM cells were permeabilized in PBS-1% BSA containing 0.1% saponin. GBM cells were stained using the Click-iT^TM^ plus Alexa Fluor^TM^ 488 picolyl azide tool kit (Thermofisher Scientific) using the manufacturer protocol. Finally, GBM cells were washed three times in PBS-1%BSA and then resuspended in 500 μl PBS-1%BSA (All steps were done at room temperature). Flow cytometry was carried out in Guava easycyte^TM^ flow cytometer. Flow data were analyzed in FlowJo software.

### RNA extraction

At the end of each experiment, 6 hydrogels per condition were combined for RNA extraction. RNA extraction was performed based on previously established protocol using Qiagen Rneasy Micro kit. Hydrogels were incubated in lysis buffer (RLT) and triturated using a 1ml syringe equipped with a 20G needle^79^. The lysate was flown through the Qiagen Qiashredder column at 17,000 G for 2 minutes. Lysates were transferred to the Rneasy micro column and RNA was extracted based on the manufacturer protocol. RNA quality was checked using a nanodrop spectrophotometer (Thermofisher).

### RNA sequencing

RNA sequencing was performed at UCLA Technology Center for Genomics & Bioinformatics (TCGB). RNA concentration and quality were checked using a Nanodrop spectrophotometer (Thermofisher). Libraries for RNA-Seq were prepared with KAPA Stranded mRNA-Seq Kit. The workflow consists of mRNA enrichment and fragmentation, first strand cDNA synthesis using random priming followed by second strand synthesis converting cDNA:RNA hybrid to double-stranded cDNA (dscDNA), and incorporates dUTP into the second cDNA strand. cDNA generation is followed by end repair to generate blunt ends, A-tailing, adaptor ligation and PCR amplification. Different adaptors were used for multiplexing samples in one lane. Sequencing was performed on an Illumina NovaSeq6000 for PE 2x150 run. Data quality check was done on Illumina SAV. Demultiplexing was performed with Illumina Bcl2fastq v2.19.1.403 software. The reads were mapped by STAR 2.27a^80^ and read counts per gene were quantified using human Ensembl GRCh38.98 GTF file. In Partek Flow, read counts were normalized by CPM +1.0E-4. All results of differential gene expression analysis utilized Partek’s statistical analysis tool, GSA. For differentially expressed gene list, *p*-values and fold change (FC) filters were applied. The filter was p<0.05 and FC>2 for all differential gene expression results. Ingenuity Pathway Analysis software (IPA, Qiagen) was used for pathway analysis. Using the list of significantly (p<0.05) differentially expressed (FC>2) genes, the Canonical Pathway analysis, Disease & Function analysis, and networks analysis were performed by IPA.

### NADH fluorescence lifetime imaging (FLIM)

NADH fluorescence lifetime images were acquired with an LSM 880 confocal microscope (Zeiss) with a 40x water-immersion objective coupled to an A320 FastFLIM acquisition system (ISS). A Ti:Sapphire laser (Spectra-Physics Mai Tai) with an 80 MHz repetition rate was used for two-photon excitation at 740 nm. The excitation signal was separated from the emission signal by a 690 nm dichroic mirror. The NADH signal was passed through a 460/80 nm bandpass filter and collected with an external photomultiplier tube (H7522P-40, Hamamatsu). Imaging was done while cell-laden hydrogels were incubated within a stage-top incubator kept at 5% CO_2_ and 37°C. FLIM data were acquired and calibrated with the SimFCS 4 software developed at the Laboratory for Fluorescence Dynamics at UC Irvine. Calibration of the system was performed by acquiring FLIM images of coumarin 6 (∼10 µM), which has a known lifetime of 2.4 ns in ethanol, to account for the instrument response function.

### Phasor FLIM NADH fractional analysis

NADH assumes two main physical states, a closed configuration when free in solution, and an open configuration when bound to an enzyme. These two physical states have differing lifetimes, 0.4 ns when in its free configuration, and 3.4 ns when in its bound configuration. To quantify metabolic alterations, we perform fractional analysis of NADH lifetime by calculating individual pixel positions on the phasor plot along the linear trajectory of purely free NADH lifetime (0.4 ns) and purely bound NADH lifetime (3.4 ns). We quantify the fraction of free NADH by simply calculating the distance of the center of mass of a spheroid’s cytoplasmic NADH FLIM pixel distribution to the position of purely bound NADH divided by the distance between purely free NADH and purely bound NADH on the phasor plot. To ensure avoiding contribution from background fluorescence which may be present in the media, we perform a three-component analysis post-background calibration to linearly unmix its contribution. We also use an empirically determined intensity threshold for each file to exclude any low-intensity background signal arising from the surrounding ECM and media. These segmentation and phasor analysis methods are described in detail elsewhere^81^.

Normalized fraction of bound NADH is calculated by normalizing the fraction of bound NADH of each sample to the median value of fraction of bound NADH in the stiff-control data set.

### Mitochondria imaging

Live cell imaging of mitochondrial structures was performed by first incubating HA-embedded spheroids in 100 nM of Tetramethylrhodamine, methyl ester (TMRM) for 1 hour at 5% CO_2_ and 37°C, then immediately placed within a stage-top incubator at the same conditions. Imaging was carried out on an LSM 880 (Zeiss) inverted laser scanning confocal microscope with a 40x, 1.2 numerical aperture, C-Apochromat water-immersion objective. A frame size of 512x512 was used, with a pixel size of 87.9 nm, and a rate of 1 frame per second for 120 frames. A two-photon titanium: sapphire laser (Spectra-Physics, MaiTai) with an 80 MHz repetition rate was used to excite the TMRM at a wavelength of 820 nm, which was passed through a 690 nm dichroic filter. The fluorescence emission in the range of 520-700 nm was captured through the microscope’s internal detector. Images were converted to TIFs in ImageJ v1.53c.

### Mitochondria structure analysis

Mitochondria content was analyzed using the mean fluorescent intensity of mitochondria structures stained with the Mitotracker green dye (Thermofisher scientific). Mean fluorescent intensity of each image was measured using ImageJ and normalized to the image area. To account for the background intensity, mean fluorescent intensity of 3 circular black areas (each with 10 μm diameter) were measured. For each image, the average value of background intensity was subtracted from the image mean fluorescent intensity to yield the final mean fluorescence intensity of the image.

Time-lapse fluorescence images of mitochondria are segmented, tracked, and analyzed using the Mitometer software. An in-depth explanation of its method can be found elsewhere^30^. Briefly, Mitometer automates the mitochondrial segmentation using a shape- and size-preserving background removal algorithm, and mitochondrial tracking by linking mitochondria from adjacent frames using both displacement differences and morphological differences followed by a gap-closing algorithm. Only mitochondrial tracks that had at least 3 track segments were kept for analysis. Mitochondria tracks were analyzed for distance as a motility feature. Distance is the summation of the linear distance across all track segments of a mitochondrial track. Additionally, mitochondria tracks were analyzed for morphology features using the solidity parameter. Solidity is the total number of pixels of the mitochondrial object divided by the number of pixels in the mitochondrion’s convex hull. Individual data points for solidity are the median values of the respective features for a single mitochondrial track.

### Metabolomics

To prepare the dried extract, 20 μl media from each hydrogel condition was transferred to pre-chilled centrifuge tubes on ice. To each tube, 500 μl chilled 80% methanol and 5 μl norvaline were added. Samples were stored at −80 °C for 20 minutes and then centrifuged at max speed for 5 minutes at 4 °C. Supernatant of each tube was transferred to a new tube and samples were dried under low pressure.

The dried extract was resuspended in 400 μl water. Using an Ion Chromatography System (ICS) 5000 (Thermo Scientific), 100 μl extract was loaded onto a Dionex IonPac AS11-HC-4μm anion-exchange column using a flow rate of 350 μl/min and separated using a 13 min gradient of 5-95 mM KOH. The attached Q Exactive mass spectrometer (Thermo Scientific) acquired full SIM data in negative polarity mode at 70K resolution with a scan range of 70-900 m/z. Metabolite data was extracted using the open-source Maven (version 8.1.27.11) software. Metabolites were identified based on accurate mass (±5 ppm) and previously established retention times of pure standards.

Quantification was done using the AreaTop feature. Data analysis was performed using in-house R scripts.

### Small molecule inhibition

Post-encapsulation, to inhibit oxidative phosphorylation and aerobic glycolysis, GBM spheroids were treated by rotenone (Sigma-Aldrich) and dichloroacetate (DCA, Sigma-Aldrich), respectively. Rotenone stock was prepared at 50mM in chloroform. DCA stock was prepared at 20mM in GBM culture media. For inhibition experiments, final concentration of 50nM rotenone and 10mM DCA were used^82,83^.

To interfere GBM cell interaction with RGD and HA, Cilengitide^TM^ (CRGD) and ezrin inhibitor was used respectively. Post-encapsulation, spheroids were treated with Cilengitide^TM^ (Sigma Aldrich) and NSC668394 (Ezrin inhibitor, Sigma Aldrich). Cilengitide^TM^ stock solution was prepared at 5 mM in PBS. NSC668394 stock solution was prepared at 10 mM in DMSO. For inhibition experiments, final concentration of 25 μM Cilengitide^TM^ and 10 μM NSC668394 were used^84,85^.

### Statistical analysis

Statistical analysis was performed in Prism 9 software. Normality of each data set was analyzed using D’Agostino & Pearson omnibus normality test. For normally distributed population, ANOVA (one or two-way) followed up by Tukey post-hoc test was performed. For non-normal distributions, Mann-Whitney non-parametric test was used.

## Author contributions

Study concept and design – AS, SKS

Acquisition of funding –AS, SKS, HIK

Acquisition of data – AS, AEYTL, MJH, MC, GS, IS, SB, NT

Analysis and interpretation of data – AS, AEYTL, MC, TMS, MJH, HIK, MAD, SK

Drafting of the manuscript – AS, SKS

Critical revision of the manuscript for important intellectual content – AS, AEYTL, HIK, MAD, SKS

## Supporting information

Supplement video 1-Soft

Supplement video 1-Stiff

