## Supplemental data and method for "Microenvironmental Stiffness Induces Metabolic Reprogramming in Glioblastoma"

**Affiliations**
1)Department of Biomedical Engineering, University of Texas at Austin
2)Department of Bioengineering, University of California, Los Angeles
3)Department of Bioengineering, University of California at Irvine
4)Department of Molecular and Medical Pharmacology, David Geffen School of Medicine, University of California, Los Angeles

Author contributions:
Study concept and design – AS, SKS
Acquisition of funding –AS, SKS, HIK
Acquisition of data – AS, AEYTL, MJH, MC, GS, IS, SB, NT
Analysis and interpretation of data – AS, AEYTL, MC, TMS, MJH, HIK, MAD, SK
Drafting of the manuscript – AS, SKS
Critical revision of the manuscript for important intellectual content – AS, AEYTL, HIK, MAD, SKS

**Supplementary Table 1**: RT-qPCR primers

| Gene | Primer name | Brand (cat. Number) |
|---|---|---|
| HYAL1 | HYAL1 TaqMan® Gene Expression Assays, Dye: FAM-MGB-small 250rxn cat#4331182 | Thermofisher (Hs00201046_m1) |
| HYAL2 | HYAL2 TaqMan® Gene Expression Assays, Dye: FAM-MGB-small 250rxn cat#4331182 | Thermofisher (Hs00186841_m1) |
| HYAL3 | HYAL3 TaqMan® Gene Expression Assays, Dye: FAM-MGB-small 250rxn cat#4331182 | Thermofisher (Hs00185910_m1) |
| MMP2 | MMP2 TaqMan® Gene Expression Assays, Dye: FAM-MGB-small 250rxn cat#4331182 | Thermofisher (Hs01548727_m1) |
| MMP9 | MMP9 TaqMan® Gene Expression Assays, Dye: FAM-MGB-small 250rxn cat#4331182 | Thermofisher (Hs00957562_m1) |
| GAPDH | GAPDH TaqMan® Gene Expression Assays, Dye: FAM-MGB-small 250rxn cat#4331182 | Thermofisher (Hs02758991_g1) |
| TGFB1 | TGFB1TaqMan® Gene Expression Assays, Dye: FAM-MGB-small 250rxn cat#4331182 | Thermofisher (Hs00998133_m1) |
| TIMP1 | TIMP1 TaqMan® Gene Expression Assays, Dye: FAM-MGB-small 250rxn cat#4331182 | Thermofisher (Hs01092512_g1) |
| CHI3L1 | CHI3L1 TaqMan® Gene Expression Assays, Dye: FAM-MGB-small 250rxn cat#4331182 | Thermofisher (Hs01072228_m1) |
| OLIG2 | OLIG2 TaqMan® Gene Expression Assays, Dye: FAM-MGB-small 250rxn cat#4331182 | Thermofisher (HS0300164_s1) |

**Supplementary Table 2**: The primary and secondary antibodies

| Antibody | Brand (Cat. Number) | Dilution |
|---|---|---|
| Rabbit monoclonal antibody to cleaved PARP | Cell Signaling Technology (5625S) | 1:200 |
| Rabbit monoclonal antibody to Ezrin | Cell Signaling Technology (3145S) | 1:200 |
| Mouse monoclonal antibody to CD44 | Cell Signaling Technology (3570S) | 1:200 |
| Mouse anti-mitochondria antibody [113-1] | Abcam (ab92924) | 1:200 |
| Alexa Fluor™ 647 Phalloidin (Actin) | Invitrogen™ (A22287) | 1:400 |
| Hoechst 33342 (Nuclei) | Thermo Scientific (62249) | 1:1000 |
| Donkey anti-mouse IgG secondary antibody, Alexa Fluor 555 | Fisher scientific (A31570) | 1:500 |
| Donkey anti-rabbit IgG secondary antibody, Alexa Fluor 647 | Fisher scientific (A31573) | 1:500 |

**Supplementary Table 3:** AFM $\mu$–compression data, collected from mice #1 and #2. Data reported as the median Young's moduli (Pa), interquartile range (Pa)

| Mice # | Core (C) | Edge (E) | Peritumoral (PT) |
|---|---|---|---|
| 1 | 3689, 2662 | 1260, 1457 | 496, 480 |
| 2 | 2781, 2338 | 1171, 733 | 263, 155 |

### Supplementary Figure 1:

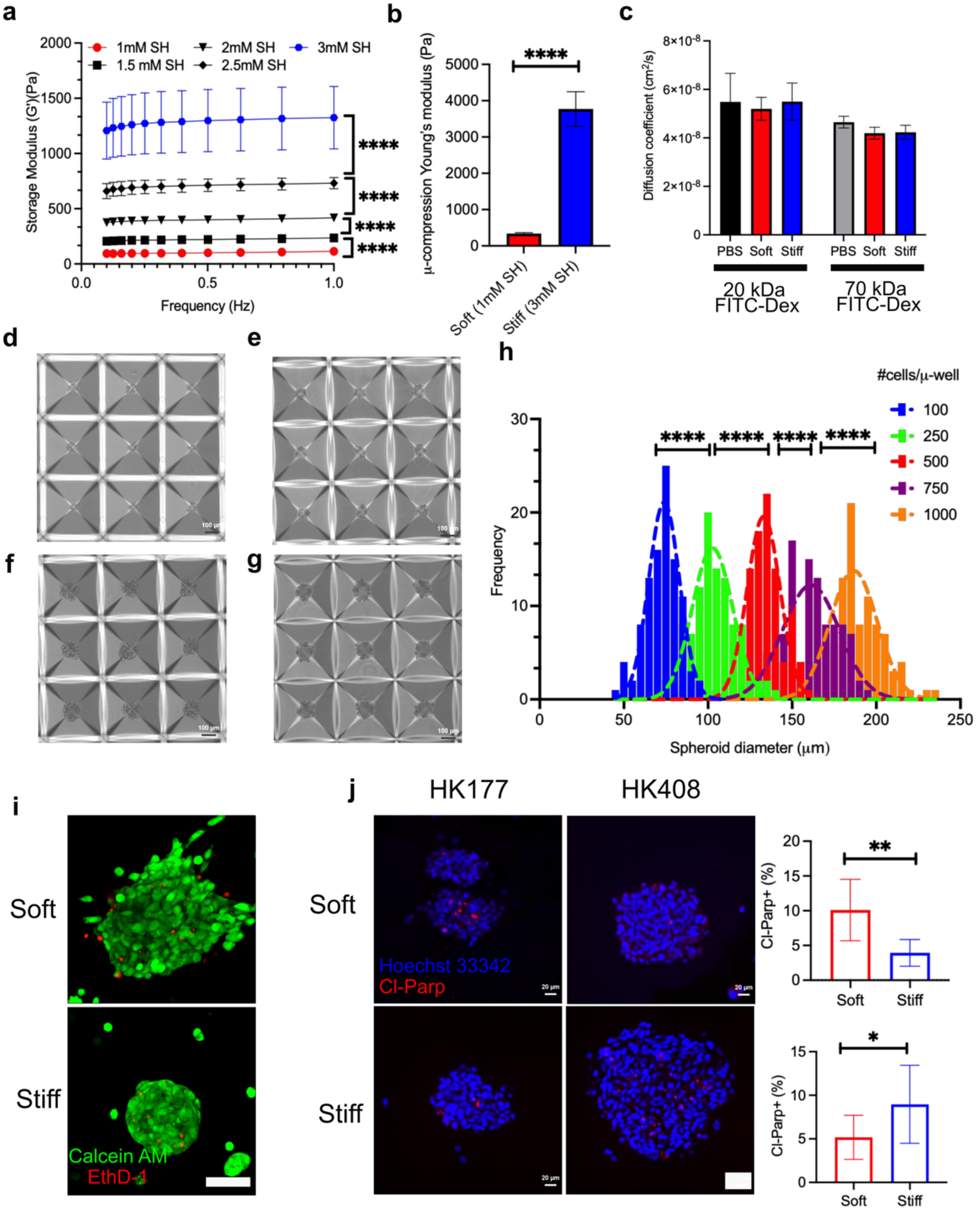


Supplementary Figure 1: 3D Hyaluronic acid hydrogels, formed via thiol-ene photochemistry. (a) By changing the total thiol content, we were able to fabricate hydrogels with a wide range of mechanical properties (n = 2 biological repeats, 5 samples measured in each biological repeat, One-way ANOVA: df=4, $p < 0.0001$, Welch's t-test: ****: $p < 0.0001$). (b) AFM measurements on hydrogels with 1mM and 3mM thiol concentrations. These hydrogels mimicked the stiffness of the GBM tumor core and the peri-tumoral brain tissue (reported values are Mean±SD Mann-Whitney $p < 0.0001$). (c) FRAP analysis of 20kDa and 70kDa dextran molecules in soft and stiff hydrogels. There were no noticeable differences in the diffusion coefficient of 20kDa and 70kDa dextran molecules in soft and stiff hydrogels. (d-h): We used Aggrewell$^{TM}$ plates to form uniform sized spheres. Sphere sizes were controlled by average seeding density. Seeding densities of (d)100 (e)250 (f)500 (g)750 cells/m-well were used (Scale bar = 100µm). (h) Sphere size distribution of different seeding densities. (Brown-Forsythe ANOVA df=4, $p < 0.0001$) (Each pair-wise test: Welch's t-test $p < 0.0001$). (i) HK408 GBM spheroids showed high survival in soft and stiff hydrogels (scale bar= 100 µm). (j) Cl-parp apoptosis marker (n = 2 biological replicates, reported values are Mean±SD Welch's t test: t= 3.621, df = 9, $p = 0.005$),(scale bar=20 µm)

**Supplementary Figure 2:**

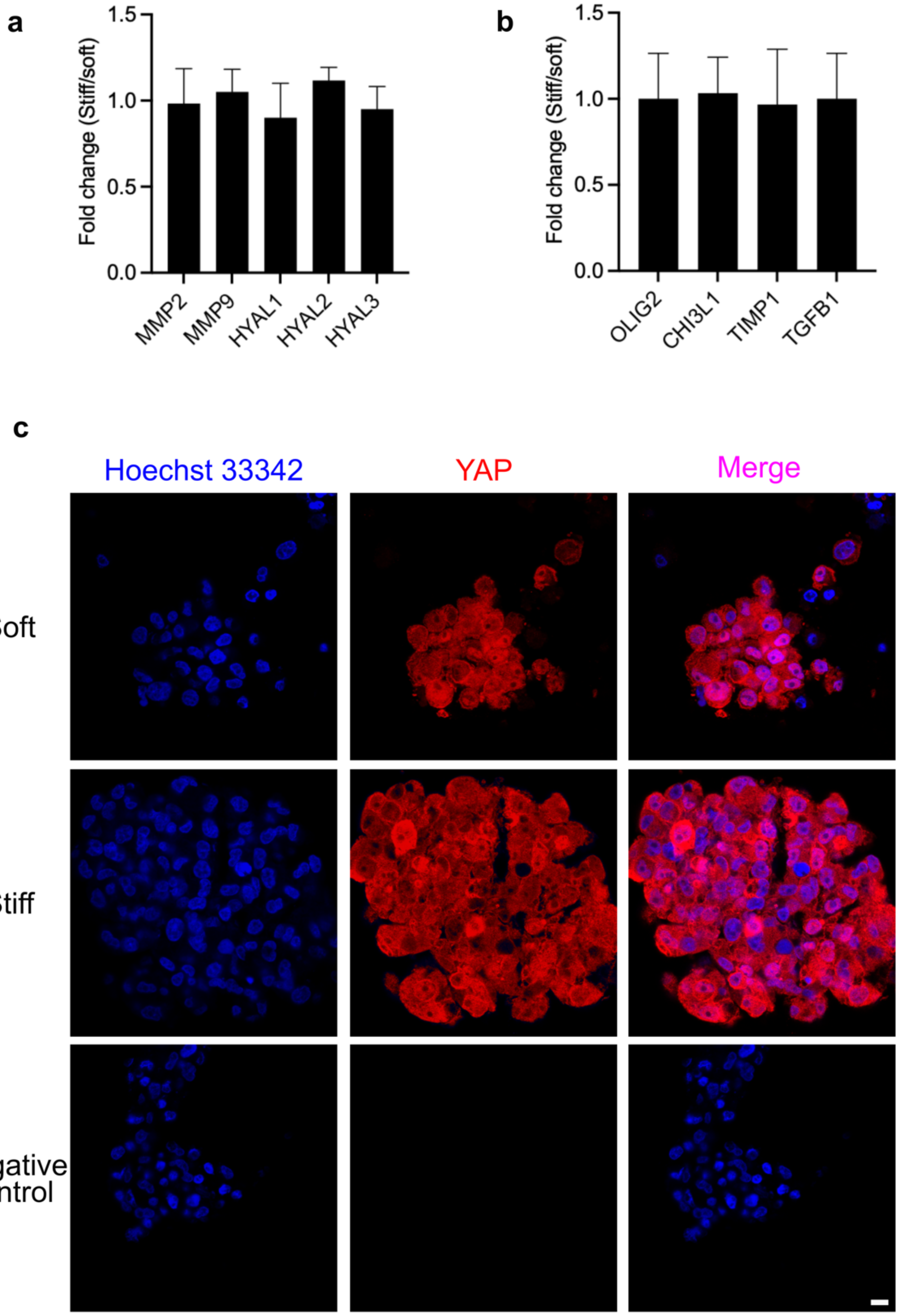


Supplementary Figure 2: In HK177 cell line RT-qPCR results of candidate genes did not show a difference in (a) matrix-degradation or (b) PMT genes in GBM cells cultured in stiff hydrogels when compared to ones cultured in soft hydrogels (reported values are Mean±SD). (c) We did not observe any changes in nuclear/cytoplasmic localization of YAP in GBM cells cultures in soft or stiff hydrogels. (Scale bar = 10μm)

**Supplementary Figure 3:**

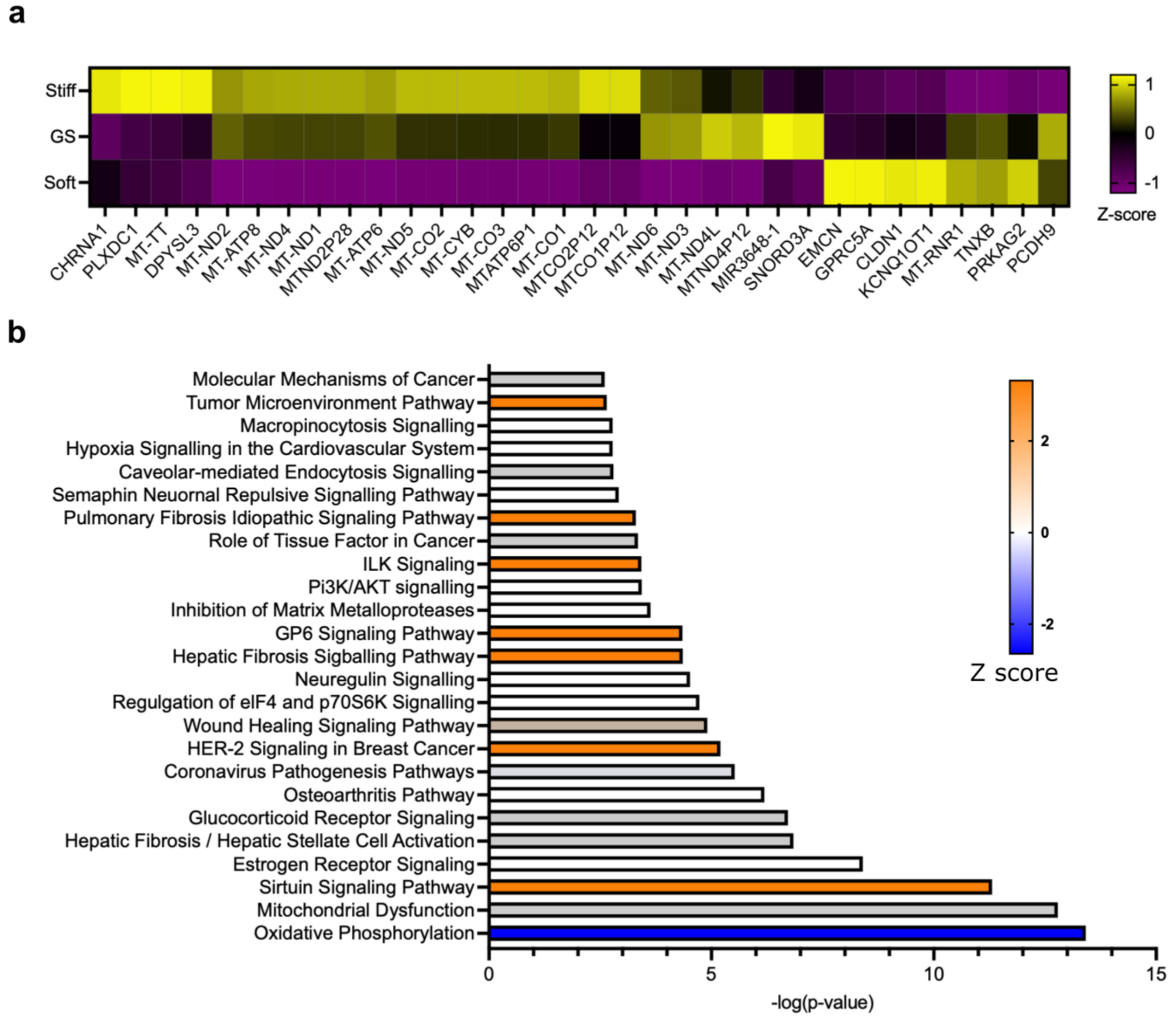


Supplementary Figure 3: (a) RNA-seq data of HK408 shows lower expression in mitochondrially-encoded genes involved in electron transport chain in soft hydrogels (b) Similar to HK177, for HK408 cell line, IPA marks oxidative phosphorylation and mitochondria dysfunction in top 3 affected canonical pathways.

**Supplementary Figure 4:**

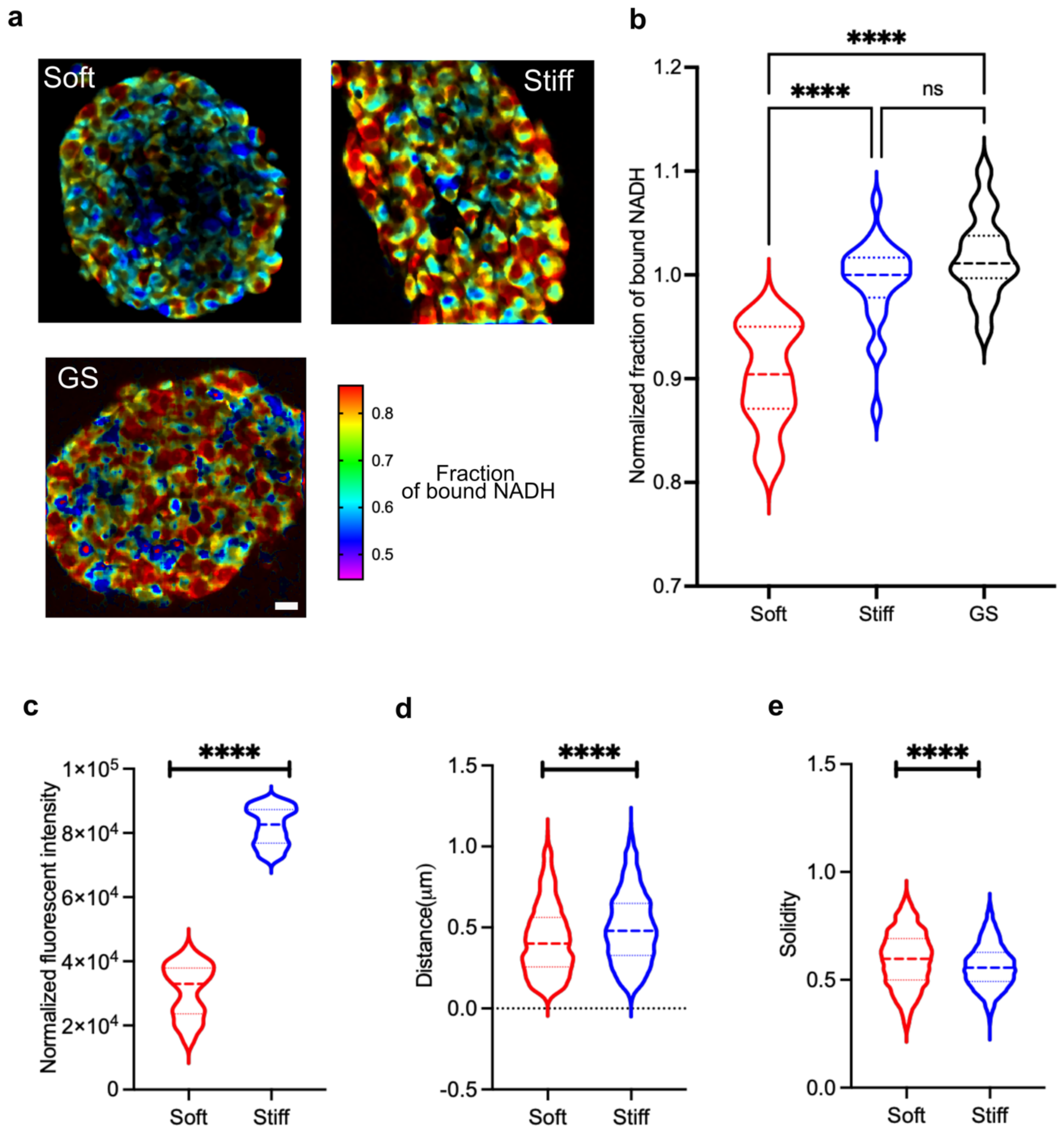


Supplementary Figure 4: (a) Representative FLIM images of HK408 (Scale bar = 20μm). (b) Lower normalized fraction of bound NADH for HK408 suggests a shift toward GLY in soft hydrogels. (Mann-Whitney $p < 0.0001$) (c) Mitochondria content (Mann-Whitney $p < 0.0001$). (d) travelled distance (Mann-Whitney $p < 0.0001$) and (e) solidity (Mann-Whitney $p < 0.0001$) all suggest higher GLY activity in soft hydrogels.

### Supplementary Figure 5

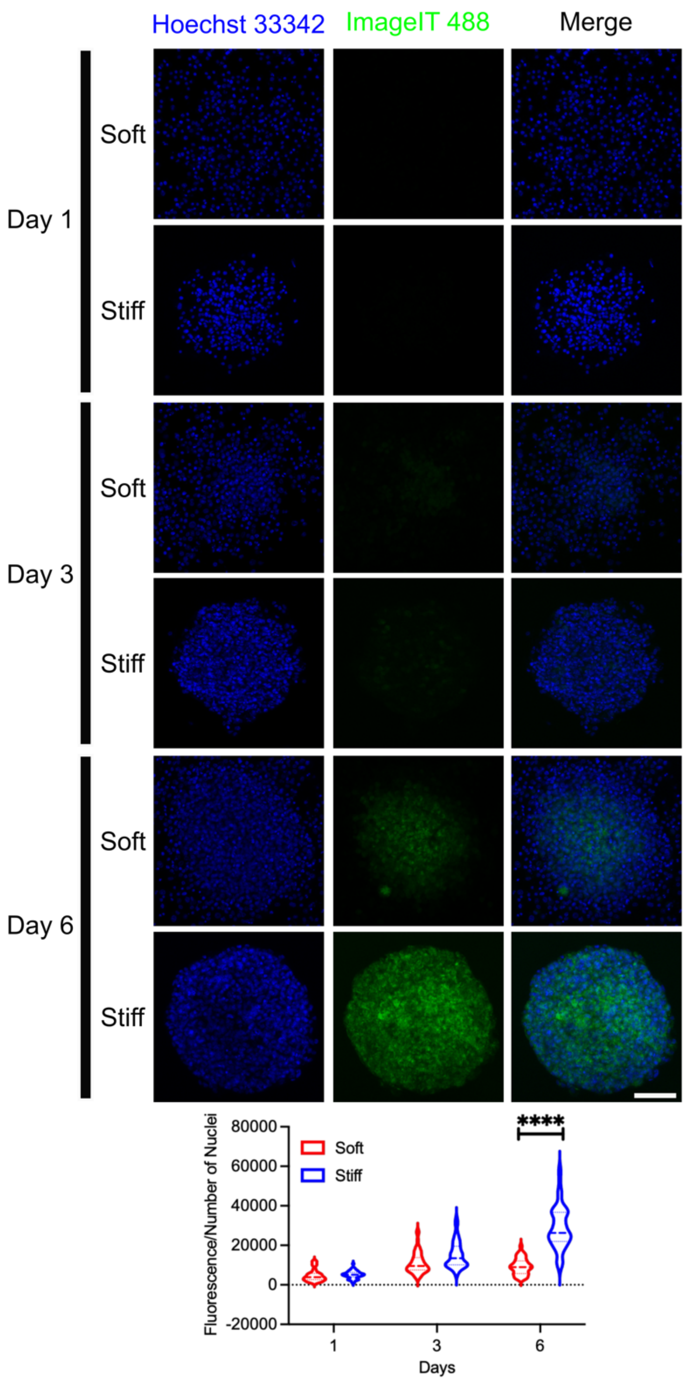


Supplementary figure 5: For HK177, oxygen availability was investigated using an Image-IT$^{TM}$ oxygen-sensitive dye. The dye becomes fluorescent at low oxygen concentrations. Interestingly, At the end of the experimental period, GBM spheroids in the center of stiff hydrogels experience lower concentration of oxygen (Scale bar = 100 μm, 2-way ANOVA test $p < 0.0001$).

**Supplementary Figure 6:**

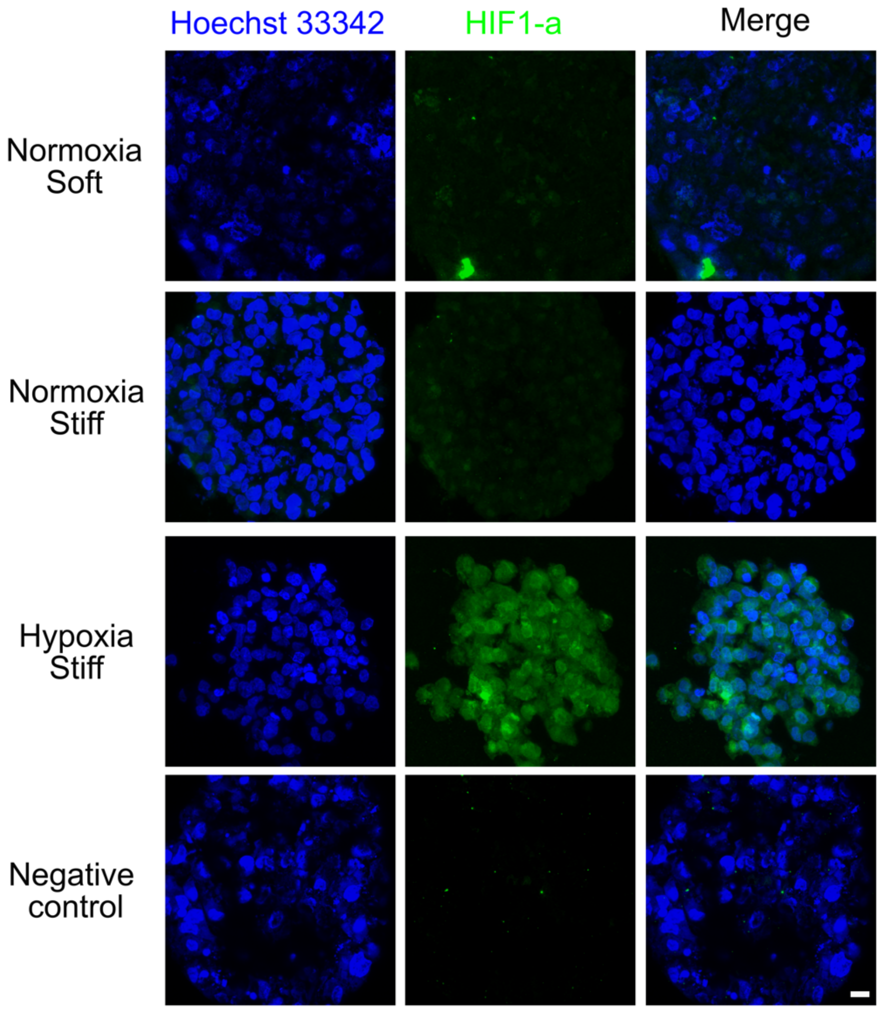


Supplementary Figure 5: Occurrence of hypoxia was investigated using immunostaining of HIF-1α in HK177 GBM spheroids. While HIF-1a was expressed in the hypoxic condition (positive control), it was expressed at a similar, low level in both soft and stiff hydrogels in normoxia. (Scale bar = 10 mm)

**Supplementary Figure 7:**

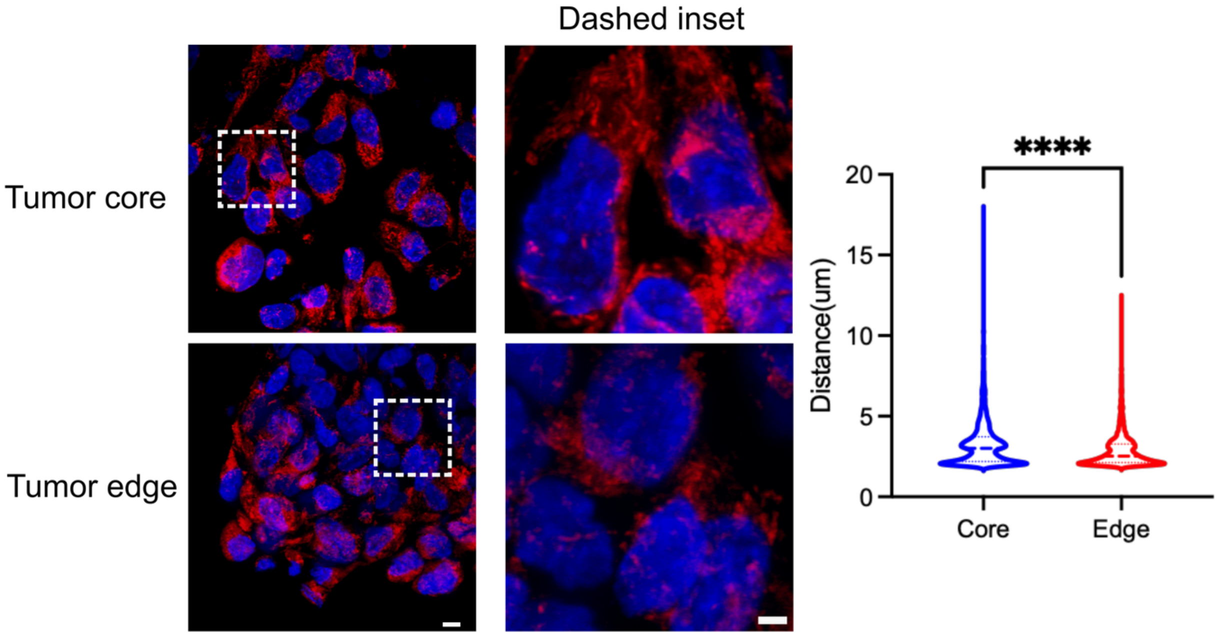


Supplementary Figure 7: Mitochondria structure was investigated in murine tumor xenografts using MT-CO1 immunostaining. Tumor cell nuclei is shown in blue (Hoechst 33342) and Mitochondria structure is shown in red (MT-CO1) (Scale bar = 10μm) (Inset scale bar = 2 μm). Mitochondria structure was analyzed using the Mitometer software. At the tumor edge, mitochondria network exhibited a smaller travelled distance which suggest a less elongated mitochondria network at the softer tumor edge when compared to the stiffer tumor core (Mann-Whitney, $p < 0.0001$).

**Supplementary Figure 8:**

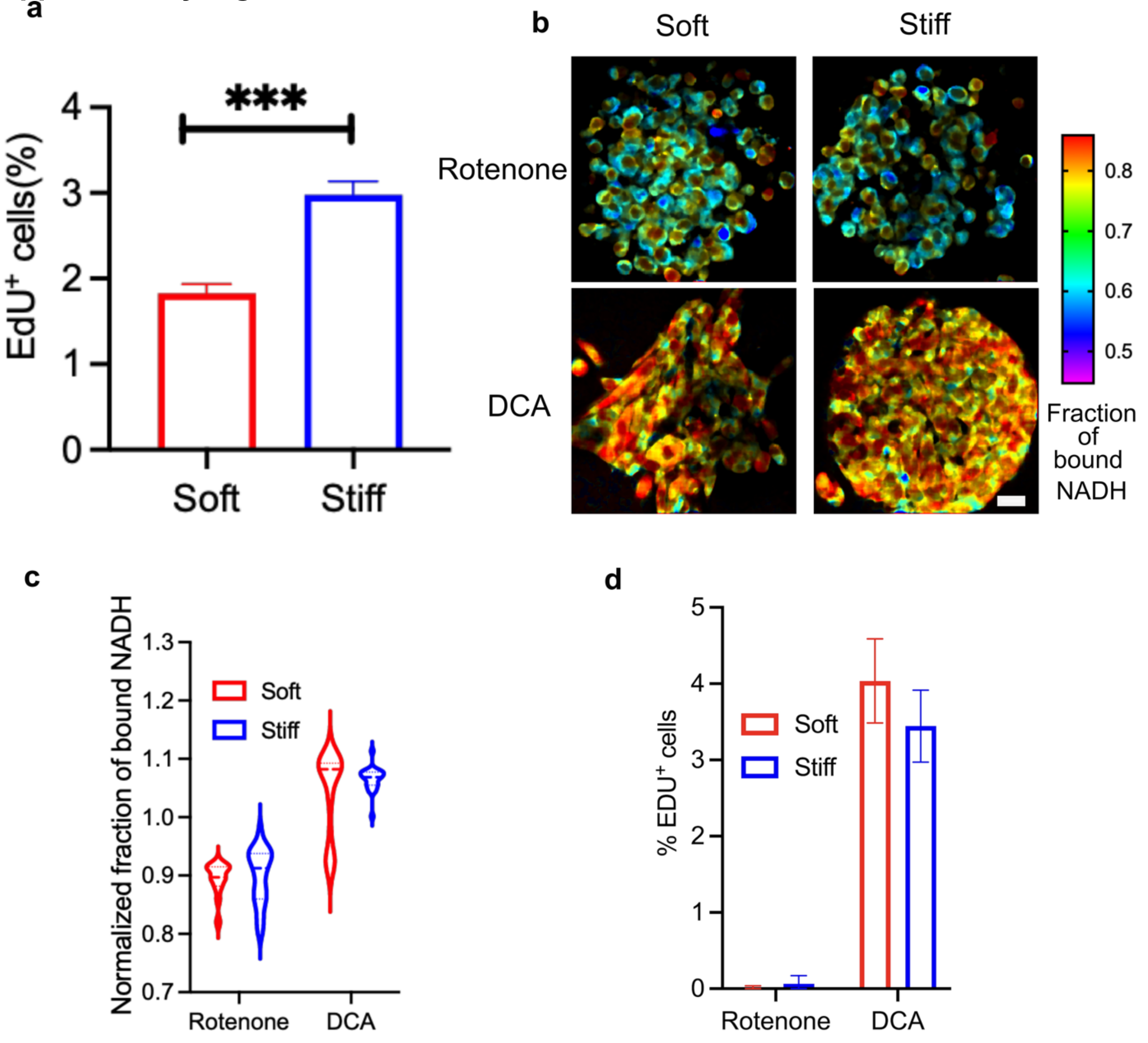


Supplementary Figure 8: (a) EdU results shows a decrease in HK408 proliferation rate in soft hydrogels n = 2 biological replicates, 3 samples per each repeat, reported values are Mean±SD, Welch's t-test $p < 0.0001$ (b)Representative FLIM images of HK408 (Scale bar = 20μm) and (c) Normalized fraction of bound NADH shows the efficacy of rotenone and DCA in altering GBM cell metabolism (2way ANOVA and Tukey post-op multiple comparison, rotenone soft vs stiff $p = 0.9954$, DCA soft vs stiff $p = 0.6379$ ). (c) EdU proliferation data (n = 2, Mean±SD, 2way ANOVA and Tukey multiple test, rotenone soft vs stiff $p = 0.9988$, DCA soft vs stiff $p = 0.2717$)

### Supplementary Figure 9:

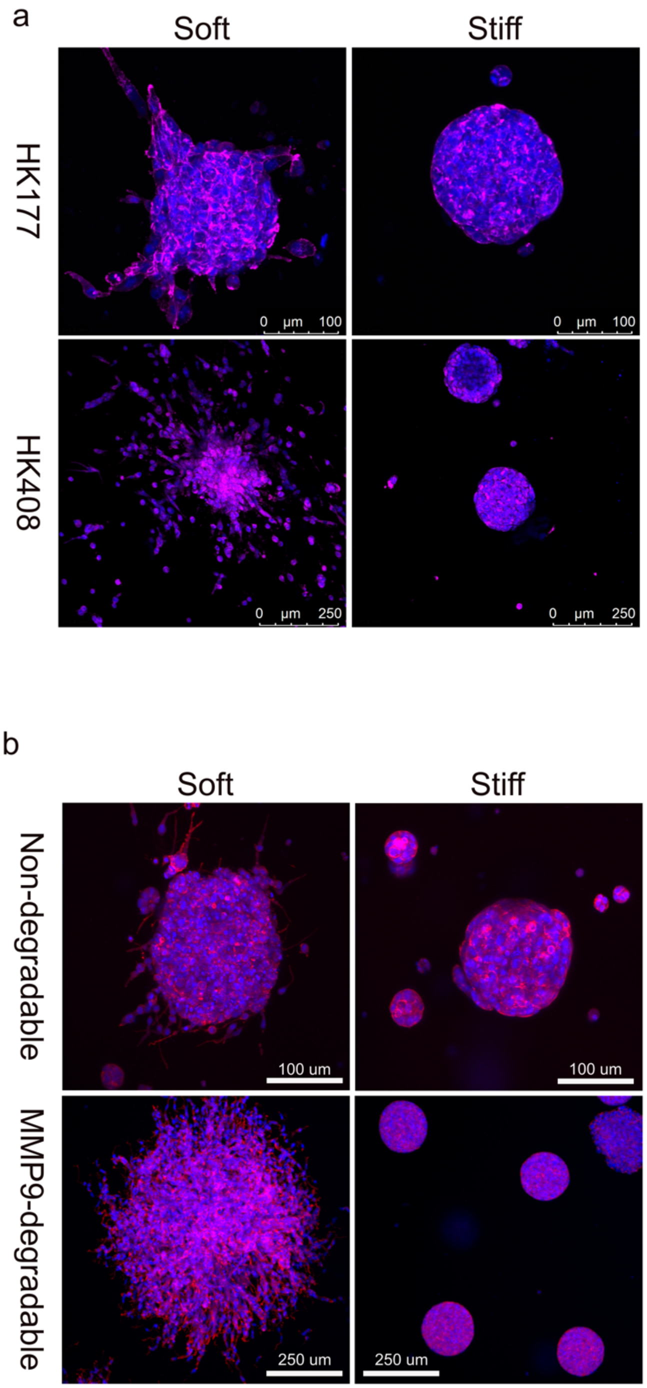


Supplementary Figure 9: (a) GBM cell migration was only observed in soft hydrogels. Migration pattern of (Scale bars = 100μm for HK177 and 250μm for HK408). (b) In order to allow enzymatic degradation of hydrogels, MMP9-degradable peptide (GCGYGVPLSLYSGYGCG) was used the cross-linker. Interestingly, inclusion of MMP9-degradable crosslinker only increased HK177 cell migration in soft hydrogels and no difference was observed in stiff hydrogels. Blue: Hoechst 33342 for nuclei. Red: Phalloidin for F-actin.

**Supplementary Figure 10:**

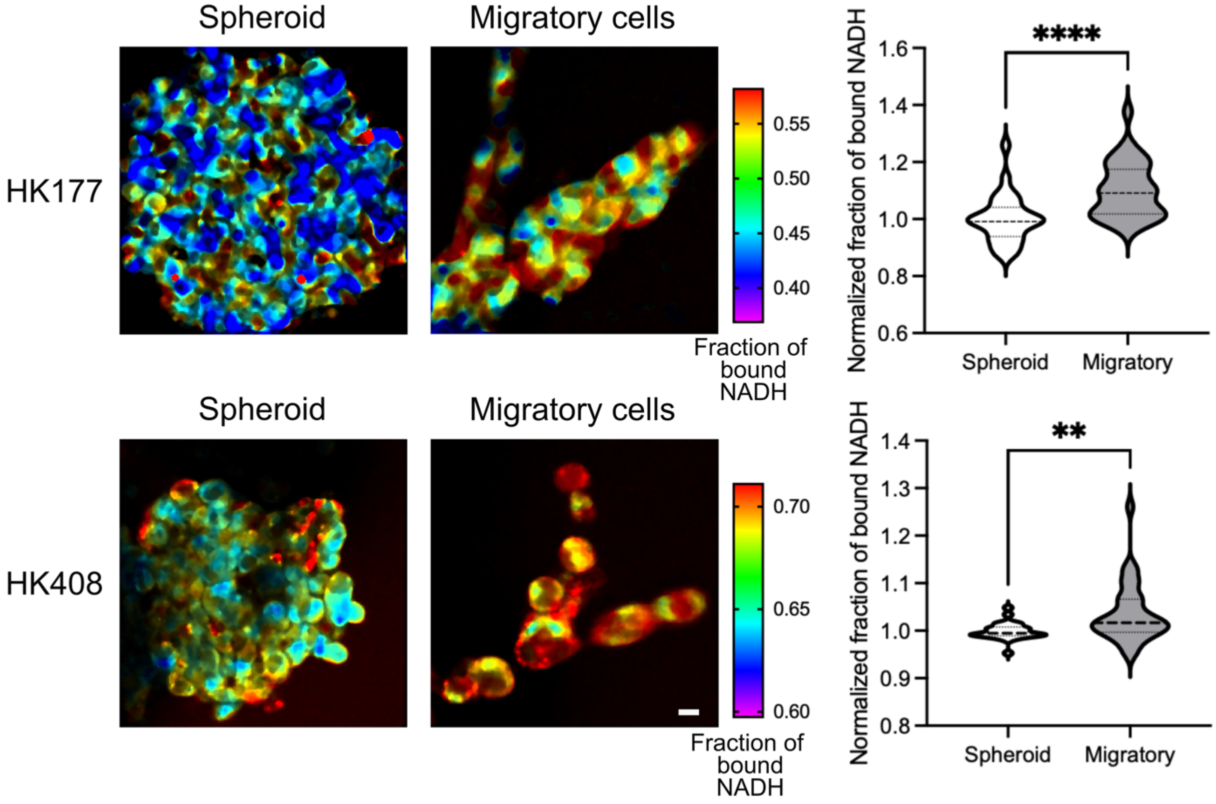


Supplementary Figure 10: FLIM data showing a shift toward OXPHOS in migratory GBM cells compared to their parental spheroid. (Scale bar = 20μm) (n=2 biological repeats with at least 5 samples in each repeat, Mann-Whitney, **: $p = 0.0036$, ****: $p < 0.0001$)

### Supplementary Figure 11:

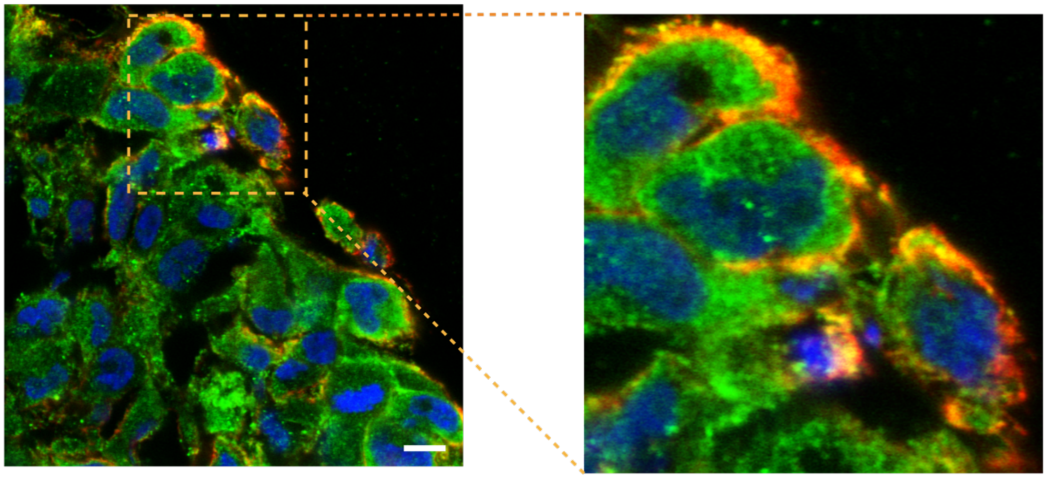

Supplementary figure 11: Immunostaining representative image of CD44(green), ezrin (red) and nuclei (blue). Close proximity of CD44 and ezrin (orange color) is visible at the periphery of GBM spheroids where GBM cells interact with HA (Scale bar = 20 μm).

### Supplementary methods:

#### HA thiolation, Hydrogel formation and characterization:

Hyaluronic acid thiolation procedure is described in our previous work[1]. Hydrogel precursor solution was prepared by dissolving HA-SH (0.5 %w/v), 4-arm thiol terminated polyethylene glycol (PEG-SH) (Laysan Bio), 8-arm norbornene terminated polyethylene glycol (PEG-Norb) (Jenkem), 0.025 %w/v lithium phenyl-2,4,6 trimethylbenzoylphosphinate (LAP, Sigma-Aldrich), 0.25 mM thiol-containing RGD (GCGYGRGDSPG, Genscript), in 20 mM HEPES buffer (pH=7). After dissolving, the hydrogel precursor solution was cast into 4mm silicone molds (Grace Biolabs) and irradiated with long-wave UV (365 nm, 4.2 mW/cm$^2$) (Blak-Ray™ B-100A UV lamp, UVP™) for 15 seconds.

Hydrogel storage moduli (G') were measured using a discovery hybrid rheometer-2 (DHR-2, TA Instruments) at 37 C. Frequency sweeps were performed under 1% constant strain in the range of 0.1 to 1.0 Hz. Storage modulus of each sample was calculated as the average value of the linear region of the storage curve from the frequency sweep plot. For statistical analysis, 3 separate measurements were taken in which 5 samples from each condition were measured. AFM measurements were done as described above.

For diffusion measurements, we used fluorescence recovery after photo-bleaching (FRAP). Hydrogels were incubated with fluorescein isothiocyanate-dextran (FITC-Dextran, 20kDa and 70kDa) solution (0.33 mg/ml in PBS) overnight. 5 pre-bleach images were taken at 10% power of 488 laser under a SP5 laser scanning confocal microscope (Leica). In order to bleach, 30 μm regions of hydrogels were exposed to a full power 488 laser (600 μm pinhole) for 20 seconds. 1000 frames of images were taken as post bleached images. $t_d$ values (time for half recovery) were calculated from fluorescence recovery graphs. Diffusion coefficients ($D_e$) were calculated using simplified Fick's law[2].

**Formation of similar-sized GBM spheroids**

As sphere size can potentially alter spheroid behavior in hydrogels[3], GBM spheroid sizes were standardized by using Aggrewell™ well plates (Stemcell Technologies) one day prior to encapsulation. To do so, Aggrewell™ wells were filled with 1 ml Pluronic F-127 solution (5 %w/v), centrifuged at 3000x G for 5 minutes and incubated for 30 minutes at room temperature. Then, Pluronic solution was aspirated, and wells were washed with 1ml complete media, cells were added, and wells were filled to 1 ml with GBM media. For all experiments, we used density of 500 cells/μ–well. Finally, the plate was centrifuged at 300x G for 3 minutes. Plates were incubated in the culture incubator overnight.

**Immunofluorescent staining**

Spheroid-laden hydrogels were fixed in 4% paraformaldehyde (PFA) for 15 minutes at 37 °C. Then, incubated with PBS solution containing 5% sucrose for 1 hour at room temperature followed by PBS solution containing 20% sucrose overnight at 4 °C. The next day, hydrogels were incubated in optimal cutting temperature solution containing 20% sucrose for 3 hours at 4 °C. Finally, blocks were frozen using dry ice+2-methylbutane mixture and stored at -80 °C. Cryosectioning was performed on a Leica cryostat 3050S to obtain 10–12 μm sections.

Immunofluorescent staining was done using an established protocol. Cryosections were air-chilled for 20 minutes, fixed with 4% PFA for 15 minutes at room temperature, washed 3X5 minutes with Tris buffer saline containing 0.1 % (v/v) Tween-20 (TBST). In case of cytoplasmic/nuclear staining, sections were permeabilized with Tris-bufferred saline containing 0.5% (v/v) triton X-100 for 15 minutes at room temperature. Blocking was carried out using a 2% (w/v) bovine serum albumin (BSA)+4% (w/v) normal donkey serum/normal goat serum (depending on the secondary) for 1 hour at room temperature. Primary antibody incubation was done in blocking solution overnight at 4 °C. The next day, sections were washed for 3X5 minutes in TBST, then incubated with secondary antibody solution (in blocking solution) for 1 hour at room temperature. Finally, sections were washed again for 3X5 minutes using TBST. Samples were imaged using a Leica Sp5 confocal microscope.

**Quantitative Real-Time PCR**

Total RNA was extracted using the RNeasy Micro Kit (Qiagen). For qPCR we used the TaqMan Fast Advanced Master Mix and the following predesigned TaqMan oligonucleotide primers and probes. The thermal cycling conditions were 20 seconds at 95 °C, followed by 40 cycles of 1 second at 95 °C and 20 seconds at 60 °C.. For each gene expression cycles were normalized to the GAPDH expression cycle. Reported values are fold change expression of stiff to soft hydrogels.

**Hypoxia assessment**

To assess oxygen availability in GBM spheroids encapsulated in soft and stiff scaffolds, hydrogels were incubated with growth media containing 5 μM Image-iT Green Hypoxia Reagent (Invitrogen) for 24 hours at 37 °C. Then, media was removed, and spheroids were incubated with fresh media containing 1:1000 Hoechst 33347 (Thermo) for 1 hour at 37 °C. GBM scaffolds were then washed with fresh media for 1 hour on a shaker at RT before being transferred to separate wells in a concavity slide (Carolina Biologics) for confocal imaging. Each hydrogel condition was imaged using a Leica SP8 Laser Confocal Microscope at 10X. Six z-stack images were taken per condition on spheroids distributed throughout the hydrogel using the same imaging parameters.

ImageJ was used for image analysis. First, the number of nuclei in each image slice was counted using a macro to ensure consistency. The nuclei (blue channel, 405 nm) images were first thresholded, then nuclei were separated using the watershed function, and finally the number of nuclei were counted using the "Analyze Particles" function and excluding particles below 50 pixels. The intensity of the hypoxia dye (green channel, 488 nm) was then measured for each image slice using by recording the integrated intensity. Then, the intensity for each slice was divided by the corresponding number of nuclei to get the average fluorescence intensity per cell for each image slice. This was repeated for each z-stack.
